# Temporal control of mitochondrial mutagenesis reveals the fate of mtDNA mutations with age

**DOI:** 10.64898/2026.09.24.754271

**Authors:** Sarah J. Shemtov, Eric Hwang, Claire S. Chung, Herbert Anson, Lucy Carrillo, Sangmin Lee, Ivetta Vorobyova, Junxiang Wan, Scott R. Kennedy, Amy R. Vandiver, Bert M. Verheijen, Whitaker Cohn, Bérénice A. Benayoun, Max A. Thorwald, Pinchas Cohen, Jonathan Wanagat, Jean-François Gout, Marc Vermulst

## Abstract

Mutations in the mitochondrial genome (mtDNA) play a critical role in the aging process and a wide variety of age-related diseases. However, it remains unclear when the mutations that drive physiological decline arise. To answer this question, we generated a new mouse model in which mitochondrial mutagenesis can be confined to a defined window of time. Surprisingly, we found that mutations that arise during the first two months of life are sufficient to drive a wide variety of age-related pathologies, and that the severity of this pathology is broadly regulated by distinct, tissue-specific selective pressures that control the fate of mtDNA mutations with age. Further, we found that selection against deleterious variants can be modulated by manipulation of mitochondrial fusion *in vitro* and *in vivo*. These observations raise the possibility that in some tissues, the pace of aging is pre-determined by events that occur early in life and that interventions targeting mitochondrial fusion may be able to slow down or reverse the expansion of these pathogenic variants. These results carry far-reaching implications for strategies aimed at preventing or delaying age-related decline.

## INTRODUCTION

MtDNA mutations play an important role in the aging process. They impair energy production, facilitate the production of reactive oxygen species and accelerate physical decline(*1*). In addition, they play a critical role in a wide variety of age-related diseases, including cancer(*2*), heart disease(*3*), diabetes(*4, 5*), and neurodegeneration(*6*). However, when the mutations that drive age-related decline arise remains unclear. Traditionally, this decline has been attributed to mutations that arise as we grow older, when pathology emerges(*7*); however, an alternative hypothesis is that age-related pathology is driven by mutations that occur early in life, but remain functionally silent for years before they promote pathology later in life. This idea is based on the multiplicity of mtDNA in human cells. Because each cell contains hundreds, or even thousands of copies of mtDNA, *de novo* mutations are initially buffered by an excess of WT genomes(*1*). A mutant genome can only compromise cellular function and contribute to cellular aging if it clonally expands enough to outnumber the WT genomes in a cell(*8*). Because this expansion process may take years, or even decades to complete, it has been suggested that the mutations that drive age-related dysfunction must arise long before pathology becomes apparent(*9*). To explore this hypothesis, we generated a mouse model that allows for temporal control of mitochondrial mutagenesis. We then confined mitochondrial mutagenesis to the first two months of life, right before the end of their developmental cycle, and tracked the mutations that arose over that time period for the remainder of the animals’ lifespan to reveal their fate and examine their impact on age-related pathology. Interestingly, various tissues exhibited differing susceptibility to early-life mutagenesis, and manipulation of mitochondrial fusion conferred resilience to an otherwise susceptible tissue *in vivo*, and in cell lines carrying deleterious mtDNA mutations through selection against the expansion of pathogenic variants. Our findings provide new insight into the timing of the aging process and reveal that the fate of mtDNA mutations is shaped by tissue-specific selection pressures that can purge deleterious mtDNA mutations from aging cells.

## RESULTS

### *Polg*^D257A-flox^ mice provide a new model to study mtDNA mutagenesis in time and space

To generate a mouse model that enables temporal control of mitochondrial mutagenesis, we genetically engineered the *Polg* gene, which encodes the mitochondrial DNA polymerase responsible for replicating the mitochondrial genome(*10*). To do so, we inserted a loxP-flanked mini-gene into intron 2 of the endogenous *Polg* locus that encodes exons 3 to 23 of the *Polg* gene (**Fig. 1A**). In addition, we added a c.770A>C mutation to exon 3 of the mini-gene to abolish the proofreading activity of the Polg protein (*Polg*^D257A-flox^) and create a well-defined “mitochondrial mutator” allele(*11, 12*). This mini-gene is seamlessly spliced into transcripts initiated from the endogenous *Polg* promoter, so that functional Polg proteins are only produced from the mini-gene. However, upon Cre-recombination the error-prone mini-gene can be excised and replaced by the endogenous WT gene (*Polg*^D257A→WT^), so that mitochondrial mutagenesis can be restricted in both time and space. For brevity, we will refer to UBC-Cre-ER^T2^ mice, unrecombined *Polg*^D257A-flox^; UBC-Cre-ER^T2^ mice and recombined *Polg*^D257A→WT^; UBC-Cre-ER^T2^ mice as WT, *Polg*^D257A-flox^ and *Polg*^D257A→WT^ mice from now on, respectively. To ensure robust recombination of the mini-gene, we tested how various Cre-promoters (ROSA26, UBC), chemicals (tamoxifen and 4-OH tamoxifen), delivery systems (chow containing 250-500mg/kg, intraperitoneal injections of 50-100mg/kg, or oral gavages of 200mg/kg), and time courses (weekly, biweekly or month-long administration depending on the delivery method) affect the efficiency of Cre-ER^T2^ based recombination in *Polg*^D257A-flox^ mice (**Fig. 1B,C; Fig. S1A-E**). These tests showed that oral gavages of tamoxifen at 200mg/kg/day, administered in two cycles of 5 consecutive days separated by a 7-day washout period, showed nearly 100% recombination at the *Polg* locus across the tissues we tested (**Fig. 1B,C**). We used this protocol to toggle the mutator allele off in all subsequent experiments.

**Figure 1.**
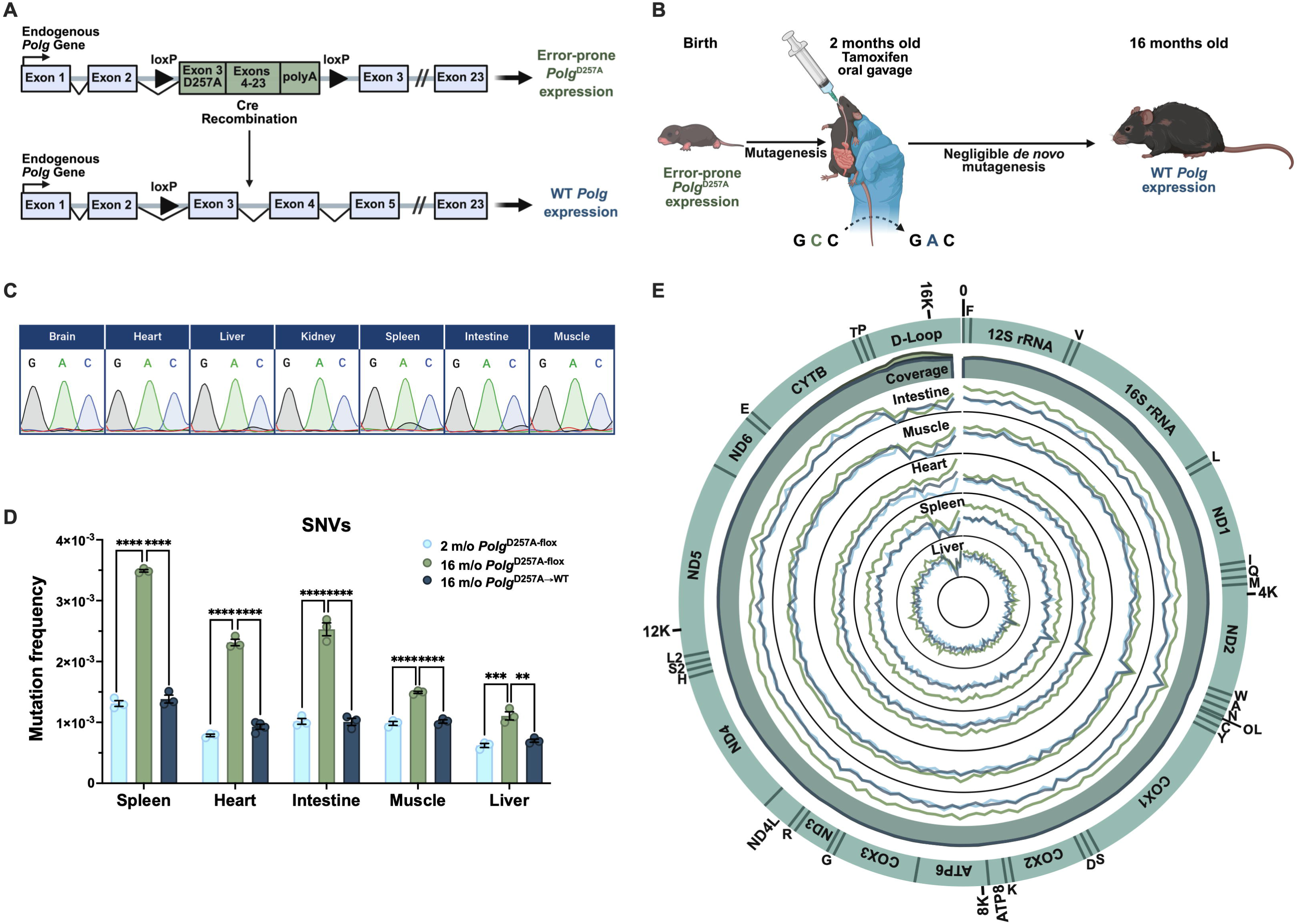
Novel mouse model and validation of experimental design. **(A)** Schematic of genetic construct that enables temporo-spatial control of mitochondrial mutagenesis (*Polg*^D257A-flox^). A minigene carrying the error-prone version of *Polg* (point mutation in exon 3, *Polg*^D257A^), encoding exons 3 to 23, followed by a termination sequence, is flanked by loxP sites and inserted within intron 2 of the endogenous *Polg* gene. Upon Cre-recombination, the minigene is excised and WT *Polg* expression is reinstated (*Polg*^D257A^ ^→WT^). **(B)** Schematic of experimental design, depicting mutant *Polg* expression until 2-months of age, when tamoxifen oral gavage enables induction of Cre Recombinase to excise the genetic construct and reinstate WT *Polg* expression for the remainder of the animals’ lifespan. **(C)** Sanger sequencing traces of cDNA across tissues (brain, heart, liver, kidney, spleen, intestine, and muscle) after Cre-Recombination, demonstrating the efficiency of the recombination process in replacing mutant *Polg* expression (*Polg*^D257A^: GCC) with WT *Polg* expression (*Polg*^D257^: GAC). **(D,E)** Mutation frequency determined by duplex sequencing of SNVs continues to rise across the spleen, heart, intestine, liver and muscle from 2-months (n=3) to 16-months in *Polg*^D257A-flox^ (n=3) animals due to ongoing mutagenesis, while the mutation burden is frozen in time in 16-month-old *Polg*^D257A→WT^ mice (n=3-4). Average coverage and mutation frequencies along the mitochondrial genome are depicted in the circle plot in **(E)**. Group differences means (± SEM) were analyzed by one-way ANOVA with Tukey’s test. *p < 0.0332, **p<0.0021, ***p<0.0002,****p<0.0001. **A** and **B** were created with BioRender.

### Recombination of the *Polg*^D257A-flox^ mini-gene freezes the mutation burden in time

To validate the *Polg*^D257A-flox^ mutator allele, as well as our strategy to control mitochondrial mutagenesis with age, we used duplex sequencing(*13*) to measure the mutation frequency of 2-to 16-month-old *Polg*^D257A-flox^ mice. Consistent with error-prone DNA synthesis by the *Polg*^D257A^ allele(*14, 15*), we found that 2-month-old *Polg*^D257A-flox^ mice exhibit a >100-fold increase in mtDNA mutation frequency compared to WT (**Fig. S2A-C**). This mutation burden was even higher in 16-month-old *Polg*^D257A-flox^ mice, demonstrating that the mutator allele continues to introduce mutations over time. In contrast, we found that if we recombined the *Polg*^D257A-flox^ allele at 2-months of age (thereby replacing it with the WT gene), further mutagenesis was prevented in all the tissues that we surveyed, including the intestine, heart, liver, spleen, and gastrocnemius muscle (**Fig. 1D, E, Fig. S2D-J)**. Thus, Cre-mediated recombination of the *Polg*^D257A-flox^ allele “freezes” the mutation burden at the time of recombination.

### Mutations that arise early in life can drive age-related pathology

Although *Polg*^D257A-flox^ mice are indistinguishable from WT mice when they are young (**Fig. S3A-D**), they display a pronounced premature aging phenotype at 16-months of age, including alopecia, inflammation, apoptosis, greying of fur, kyphosis, decreased body weight, loss of subcutaneous fat and atrophy of the testes (**Fig. 2, Movie S1**). Importantly, these symptoms closely match the natural aging phenotype of humans, as well as the phenotype of the unconditional *Polg*^D257A^ mouse model that our temporal strategy is based on(*11, 12*). To test when the mutations that cause these symptoms arise, we recombined the *Polg*^D257A-flox^ allele at 2-months of age to halt mitochondrial mutagenesis, and aged the resulting *Polg*^D257A→WT^ mice to 16-months of age as well. Interestingly, we found that *Polg*^D257A→WT^ mice developed a similar premature-aging phenotype as unrecombined *Polg*^D257A-flox^ mice, including reduced body weight (**Fig. 2E**), loss of fat mass compared to lean mass (**Fig. 2F,G**), testicular atrophy (**Fig. 2H,I**), alopecia and graying of fur (**Fig. 2D**). These age-related changes are visibly apparent in **Movie S1**, where *Polg*^D257A-flox^ (green arrow) and *Polg*^D257A→WT^ (blue arrow) look significantly more aged than their WT littermates (other two cagemates) at 16-months of age. In addition, aged (but not young, **Fig. S3E,F**) *Polg*^D257A→WT^ mice displayed reduced grip strength (p<0.0001, **Fig. 2J**) and endurance (p=0.003, **Fig. 2K**), as well as increased apoptosis (p=0.023 and p=0.018) and inflammation (p=0.0016 and p<0.0001) in liver and heart tissue respectively (**Fig. 2L-P**), with no differences in proliferation among the groups (**Fig. S4**). Finally, *Polg*^D257A→WT^ also displayed elevated plasma levels of GDF15 (p=0.04, **Fig. 2Q**), a marker of mitochondrial stress in aging humans(*16*). Because mitochondrial mutagenesis was restricted to the first two months of life, these findings demonstrate that mutations acquired during this exceptionally brief, early-life window are sufficient to drive pathology more than a year later.

**Figure 2.**
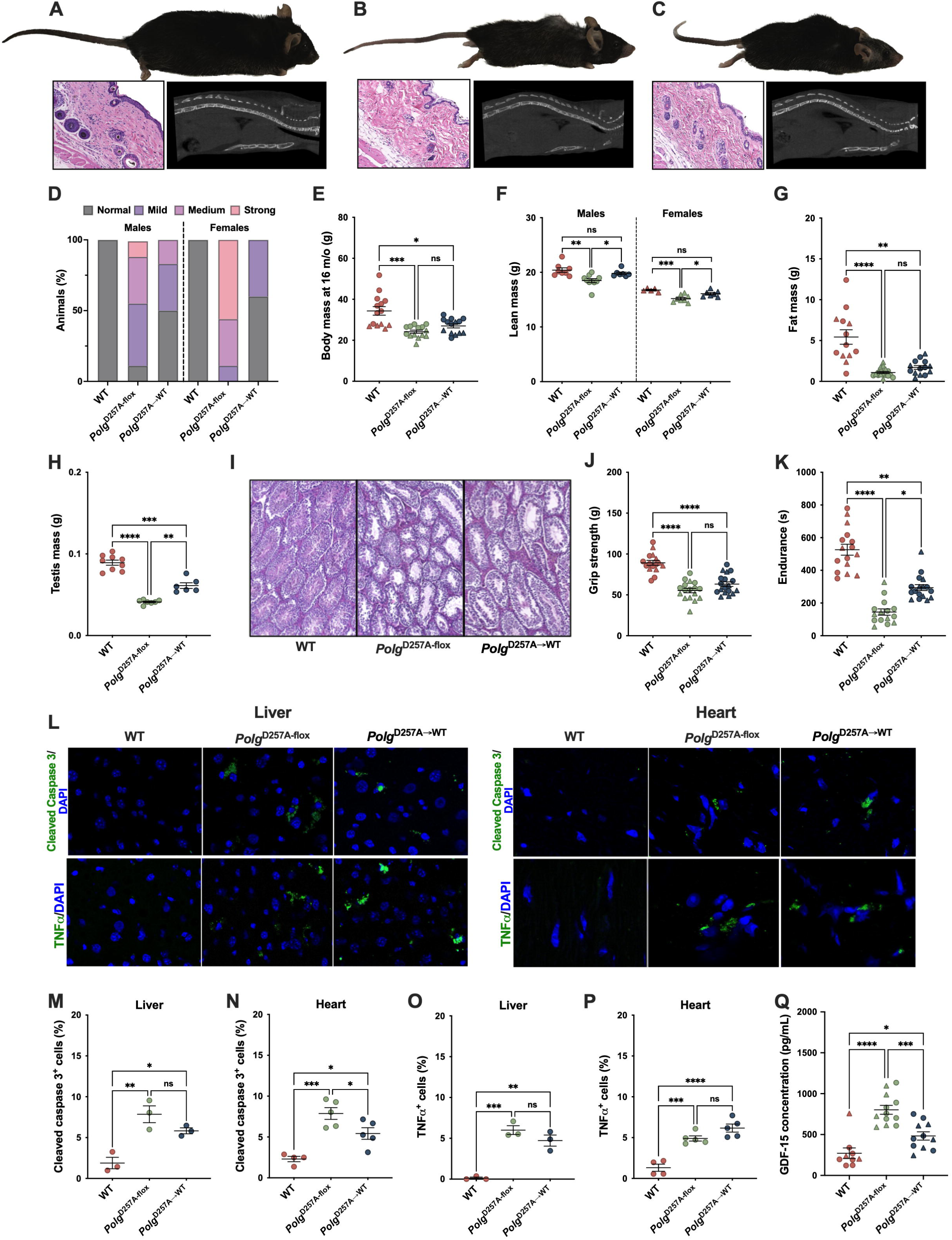
Early-life mtDNA mutations result in numerous accelerated aging phenotypes of *Polg*^D257A-flox^ mice. (A-C) Images of WT **(A)**, *Polg*^D257A-flox^ **(B)**, *Polg*^D257A→WT^ **(C)** animals at 16-months of age, along with their corresponding spinal morphology and full thickness skin H&E staining (below each animal). **(D)** Proportion of experimental animals with normal coat, or mild, medium or strong fur graying/alopecia (male n=5-9/group; female n=4-9/group). **(E)** Body weights at 16-months (male n=8-9/group; female n=5-7/group). **(F,G)** Body composition at 16 months, with lean mass depicted in **(F)** and fat mass in **(G;** male n=7-10/group; female n=5-7/group). **(H,I)** Testes are smaller **(H)** and present with altered morphology **(I)** in both *Polg*^D257A-flox^ and *Polg*^D257A→WT^ animals compared to WT (n=6-9/group). **(J,K)** Grip strength **(J)** and endurance **(K)** are significantly reduced compared to WT in both *Polg*^D257A-flox^ and *Polg*^D257A→WT^ mice. **(L-P)** Immunofluorescence **(L)** in WT, *Polg*^D257A-flox^, and *Polg*^D257A→WT^ males, with quantification, of cleaved-caspase 3 (liver, **M** and heart **N**) and TNFα (liver, **O** and heart, **P**). Images at 63x magnification; n=3-5/group. **(Q)** GDF-15 levels are elevated in plasma of *Polg*^D257A-flox^ and *Polg*^D257A→WT^ animals compared to WT (male n=6-7/group; female n=3-5/group). Group differences means (± SEM) were analyzed by one-way ANOVA with Tukey’s test, Kruskal-Wallis with Dunn’s test for nonparametric distributions, or Welch’s ANOVA when variances were unequal across groups. *p < 0.0332, **p<0.0021, ***p<0.0002,****p<0.0001. Males are denoted by: , while females: .

### Several tissues are resilient to early-life mutations

Although many of the symptoms of premature aging in *Polg*^D257A-flox^ mice are recapitulated in *Polg*^D257A→WT^ mice, some were notably absent. For example, old (but not young, **Fig. S5A-I**) *Polg*^D257A-flox^ mice display a significant increase in both heart size and weight (p=0.0002, **Fig. 3A,B**), a common symptom of cardiomyopathy in aging humans as well. This increase is accompanied by a larger internal diameter of the left ventricle in systole (p=0.0003) and diastole (p=0.0282) (**Fig. 3C, Fig S5J**), as well as reduced ejection fraction (p<0.0001, **Fig. 3D**) and fractional shortening (p<0.0001, **Fig. 3E**). *Polg*^D257A-flox^ mice also exhibit an altered E/A ratio (p=0.0002) that is primarily driven by reduced A-wave velocity (p=0.0022). These measurements provide evidence of widespread structural and functional decline in *Polg*^D257A-flox^ hearts (**Fig. S5K-M**). Surprisingly though, cardiac pathology was almost completely absent in the hearts of *Polg*^D257A→WT^ mice (**Fig. 3A-E, Fig. S5J-M**). Similarly, old (but not young**, Fig. S5N,O**), *Polg*^D257A-flox^ mice exhibit enlarged spleens with disorganized tissue morphology (**Fig. 3F-H**) and increased plasma levels of the inflammatory marker IL-6 (**Fig. 3I**). Both of these phenotypes were absent in *Polg*^D257A→WT^ mice (p<0.0001 for spleen size and p=0.0046 for IL-6, both compared to *Polg*^D257A-flox^; **Fig. 3F-I**). Thus, although early-life mtDNA mutations are sufficient to drive a wide array of age-related pathologies, their long-term consequences differ markedly between tissues, implying the existence of tissue-specific mechanisms that determine the fate of mtDNA mutations over time.

**Figure 3.**
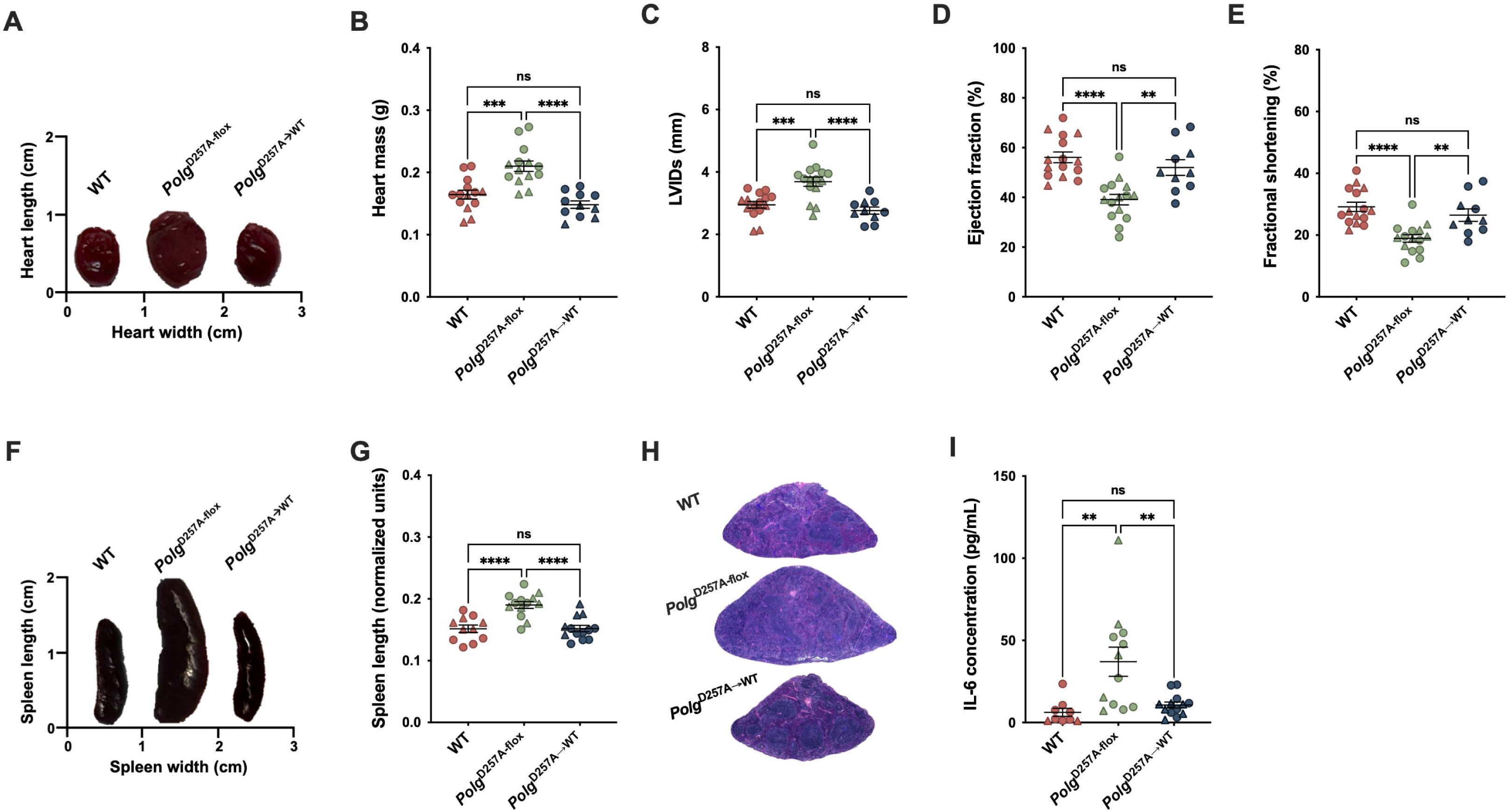
Heart and spleen are protected against early-life mtDNA mutations. (A,B) Hearts of WT and *Polg*^D257A→WT^ mice are significantly smaller in size **(A)** and mass **(B)** than those of *Polg*^D257A-flox^ animals at 16-months of age (male n=6-9/group; female n=5-7/group). **(C-E)** Heart function, specifically determined by LVIDs **(C)**, ejection fraction **(D)**, and fractional shortening **(E)**, is also unaffected in 16-month-old *Polg*^D257A→WT^ compared to *Polg*^D257A-flox^ animals (male n=6-9/group; female n=4-7/group). **(F,G)** Spleens of WT and *Polg*^D257A→WT^ mice are significantly smaller in overall size and length than those of *Polg*^D257A-flox^ animals at 16-months of age (male n=6-8/group; female n=5-6/group). **(H)** Overall spleen morphology in *Polg*^D257A→WT^ mice is also restored compared to *Polg*^D257A-flox^ animals. **(I)** IL-6 levels are elevated in plasma of *Polg*^D257A-flox^ compared to WT but are restored in *Polg*^D257A→WT^ (male n=6-7/group; female n=3-6/group). Group differences means (± SEM) were analyzed by one-way ANOVA with Tukey’s test. *p < 0.0332, **p<0.0021, ***p<0.0002,****p<0.0001. Males are denoted by: , while females: .

### Cardiac resilience extends to the molecular level

To explore the extent to which the heart is resilient to early life mutations, we examined the transcriptome, proteome and metabolome of WT, *Polg*^D257A-flox^ and *Polg*^D257A→WT^ hearts. In total, we found that 608 transcripts and 954 proteins were deregulated (p_adj_<0.05) in prematurely aged *Polg*^D257A-flox^ hearts compared to WT hearts (**Fig. 4A,B**). Together, these molecules provide evidence of extensive cardiac remodeling and metabolic alterations, including widespread depletion of respiratory-chain proteins (**Fig. 4C, Fig. S6A**). For example, proteomic analyses indicated substantially altered one-carbon and amino acid metabolism, increased hypertrophic signaling (Myh7 (protein and transcript), Ankrd1, and *Nppb)*, and modified calcium handling (Atp2a2/SERCA2, Ryr2, Mcu, Micu1, and Mcur1; **Fig. 4A-D**). In addition, 43 metabolites were significantly deregulated in *Polg*^D257A-flox^ hearts, many of which are controlled by the proteins that were significantly altered (**Fig. 4E,F**). For example, Shmt2 (p_adj_=7.7×10^−6^), Mthfd2 (p_adj_=8.2×10^−9^), Phgdh (p_adj_=1.7×10^−6^), Psat1(p_adj_=3.2x 10^−6^) are all part of the tightly interconnected serine biosynthesis and mitochondrial one-carbon metabolism pathways(*17*). Accordingly, *Polg*^D257A-flox^ hearts showed elevated serine, sarcosine and N-formylmethionine, together with the transsulfuration intermediates methionine and cystathionine. Similarly, other significantly altered proteins, like Pycr2, Dhodh, and Bckdh (subunits a and b), directly contribute to the observed deregulation of metabolites like proline, orotic acid and branched chain amino acids, respectively. *Polg*^D257A-flox^ hearts also accumulated the β-oxidation intermediate hydroxyhexanoylcarnitine and the ketone body β-hydroxybutyrate, suggesting impaired fatty acid and ketone oxidation. All these pathways are either implicated in, or respond to mitochondrial dysfunction, oxidative stress, or heart failure (**Fig. 4E**). Remarkably though, only 22 proteins and 23 transcripts were differentially expressed between *Polg*^D257A→WT^ and WT hearts (**Fig. 4A,B**), while the number of significantly deregulated metabolites was only 16, greatly preventing the deregulation of these metabolic pathways (**Fig. 4E,F**). For example, over-representation analysis (ORA) of the transcriptomics revealed that many of the genes in cluster 2 (**Fig. S6B,** by LRT), which were downregulated in *Polg*^D257A-flox^ hearts compared to WT and normalized in *Polg*^D257A→WT^, were involved in muscle contraction (p_adj_=1.22×10^−6^) and ion transport (p_adj_=3.70×10^−5^). While *Gdf15* transcripts were significantly reduced in *Polg*^D257A→WT^ mice compared to *Polg*^D257A-flox^, they remained significantly upregulated compared to WT (**Fig. S6C**), just as we observed in the plasma of these animals (**Fig. 2Q**). Together, these findings demonstrate that the physiological resilience of the heart is mirrored by near-complete molecular recovery, preventing the pathological remodeling that characterizes *Polg*^D257A-flox^ hearts (**Fig. 4D**).

**Figure 4.**
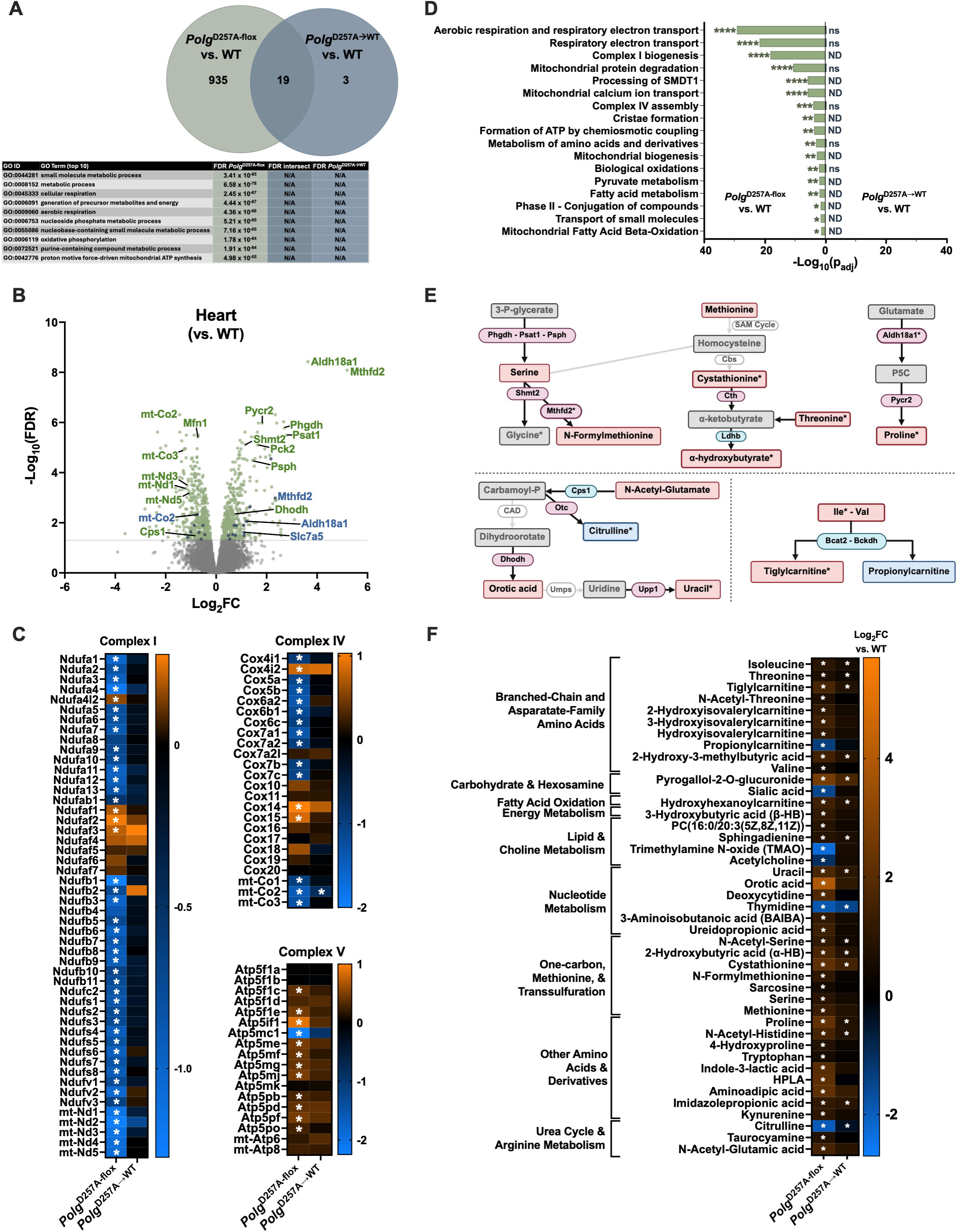
Restricting mutagenesis to early-life alleviates the disease-associated proteome and metabolome in the heart of male *Polg*^D257A→WT^ mice. (A) Unique and overlapping differentially expressed proteins between *Polg*^D257A-flox^ or *Polg*^D257A→WT^ and WT in heart. Top 10 Gene Ontology (biological process) terms are displayed in the table for significantly altered proteins. GO enrichment analysis was performed using the GO Knowledgebase, and pathways with p_adj_<0.05 were considered significant. None were identified in the intersect nor in the *Polg*^D257A→WT^ vs. WT contrasts. (B) Volcano plot of differentially expressed proteins in the heart between *Polg*^D257A-flox^ and WT (green) or *Polg*^D257A→WT^ and WT (blue). Select proteins are labeled. (C) Heatmap of differential protein expression in *Polg*^D257A-flox^ and *Polg*^D257A→WT^ hearts compared to WT of nuclear- and mitochondrial-encoded ETC Complexes. Asterisks denote significantly altered proteins versus WT, Benjamini-Hochberg (BH) p_adj_<0.05. **(D)** ORA pathways significantly enriched in *Polg*^D257A-flox^ and *Polg*^D257A→WT^ hearts compared to WT, plotted by significance (−log_10_p_adj_). **(E)** Coordinated alterations in enzyme-metabolite pathways in *Polg*^D257A-flox^ vs. WT are dramatically dampened in *Polg*^D257A→WT^ vs. WT (significance denoted by asterisks). Colors indicate log_2_FC for *Polg*^D257A-flox^ vs. WT, with red (positive) and blue (negative). Gray denotes not significant or not detected. Created with BioRender. **(F)** Heatmap of differential metabolite abundance between *Polg*^D257A-flox^ or *Polg*^D257A→WT^ and WT. Proteomics n=6/ genotype, significance by BH p_adj_<0.05; metabolomics n=6/genotype, significance by one-way ANOVA with Tukey’s post-hoc test p_adj_<0.05.

### Clonal expansion of mtDNA mutations differs across tissues of *Polg*^D257A→WT^ mice

Because mtDNA mutations only become pathogenic after clonal expansion, tissues could protect themselves from mutations by preventing this process. To test whether this mechanism underlies the resilience of heart and spleen tissue to early-life mutations, we stained tissues with NBTx, a histochemical reagent that labels cells in which clonally expanded mtDNA mutations impair oxidative phosphorylation(*18*). NBTx is the ideal tool for these experiments, because it not only reveals the presence of clonally expanded mutations, but also their physiological consequences(*18*). At 2-months of age, tissues are devoid of NBTx positive cells, as mutations have not yet clonally expanded (**Fig. S7**). However, at 16 months, we found that unrecombined *Polg*^D257A-flox^ mice harbor large numbers of NBTx-positive cells in all the tissues we monitored, including the heart and spleen (**Fig. 5A-E**). In contrast though, we found that heart, spleen and muscle of *Polg*^D257A→WT^ mice harbored significantly fewer NBTx-positive cells compared to *Polg*^D257A-flox^ mice (p=0.0007, p=0.0156 and p<0.0001, respectively, **Fig. 5C-E**). Meanwhile, the liver (p<0.0001) and intestine (p<0.0001) of *Polg*^D257A-flox^ and *Polg*^D257A→WT^ mice displayed a similar increase in NBTx-positive cells compared to WT (**Fig. 5A,B**) , indicating that these tissues fail to prevent clonal expansion even when mutagenesis is halted. This dysfunction was reflected at the molecular level as well. Even though *Polg*^D257A→WT^ livers display certain normalized molecular patterns at the protein (**Fig. S8A**) and RNA level (**Fig. S8B**), particularly those pertaining to mitochondrial translation (**Fig. S8C**), they retained persistent, widespread evidence of mitochondrial dysfunction, including 132 differentially expressed proteins and 33 altered metabolites, many of which were shared with unrecombined *Polg*^D257A-flox^ mice (**Fig. S8A,C-F**). Overall, these molecules carried signatures of inflammatory and metabolic stress pathways, as well as respiratory-chain and tricarboxylic acid-cycle remodeling, indicating that unlike heart tissue, early-life mtDNA mutations exert profound molecular consequences in the liver. For example, complex IV proteins and assembly factors were particularly impacted in *Polg*^D257A→WT^ livers compared to WT (**Fig. S8C,D**), a readout directly captured by NBTx staining. Because heart, spleen, muscle, liver and intestine harbor comparable amounts of mtDNA mutations at the time of recombination, these results suggest that the fate of those mutations differs substantially from tissue to tissue.

**Figure 5.**
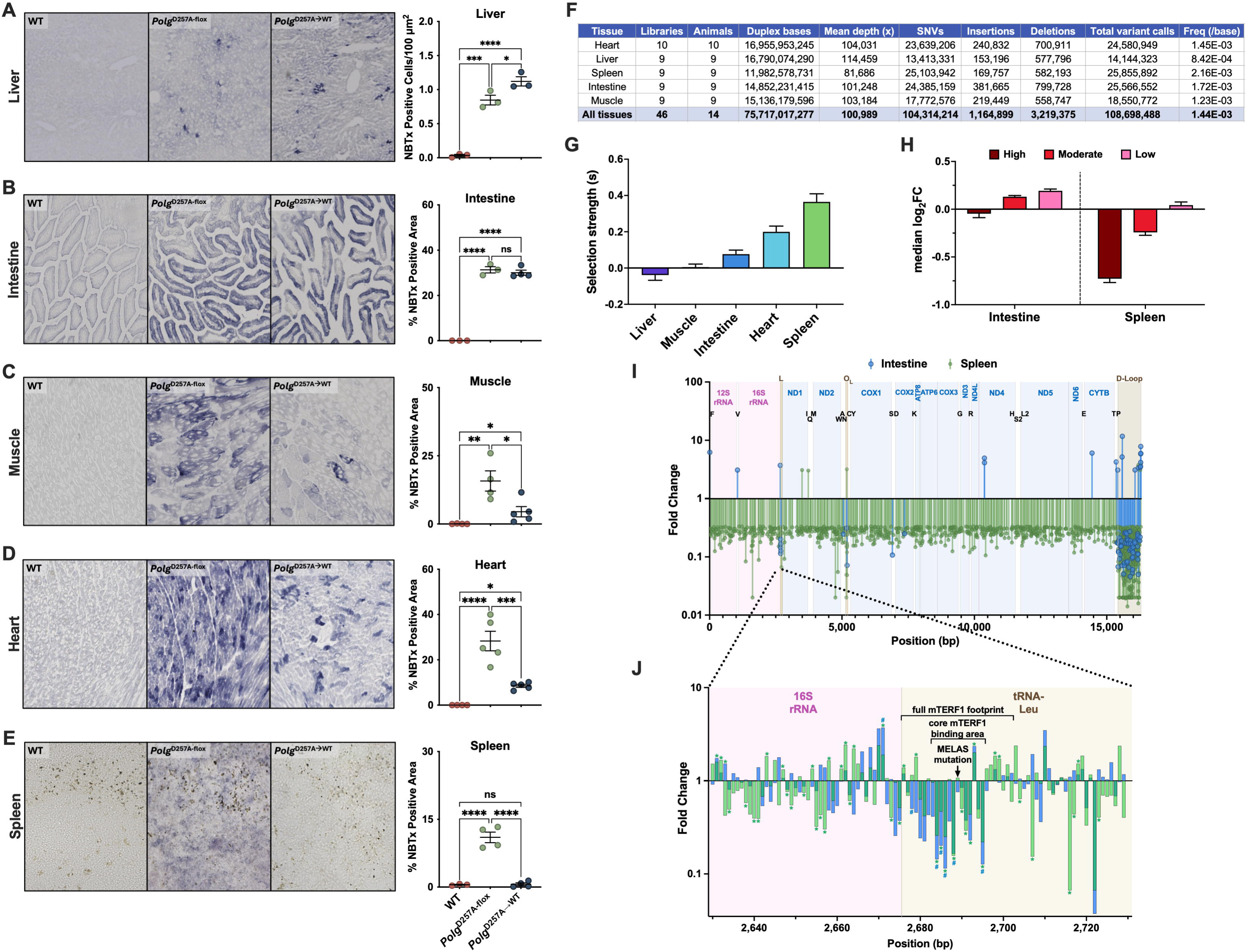
Clonal expansion of early-life mtDNA mutations and selection are tissue specific. (A-E) NBTx staining and quantification of liver **(A)**, intestine **(B)**, gastrocnemius muscle **(C)**, heart **(D)** and spleen **(E)**. Images at 10x magnification; n=3-5/group. **(F)** Table characterizing the size and depth of the duplex sequencing tissue datasets analyzed in this study. **(G)** Trajectory-based selection coefficients were estimated for each tissue as the difference between median log_2_FC of synonymous and non-synonymous variants from duplex sequencing of mtDNA. Bars represent selection coefficients, with error bars denoting 95% bootstrap CIs. Positive values indicate purifying selection acting against non-synonymous variants, with selection strength varying across tissues. **(H)** Median log_2_FC in frequencies of mtDNA mutations between 16-month- and 2-month-old *Polg*^D257A→WT^ mice of high, moderate, and low impact variants in intestine and spleen. Bars indicate medians across variants within tissue and impact class, with error bars representing 95% confidence intervals (CI) estimated by non-parametric bootstrap resampling. **(I)** Significant absolute >3-fold changes in mutation frequency over time in spleen (green) and intestine (blue). Nearly all significant >3-fold changes occur in regulatory regions in the intestine, while selection is consistent across the genome in spleen. **(J)** Inset of **I**, focused on the mTerf1 binding region in mouse mtDNA. Each bar represents the fold change of mutations at a single mtDNA base in either spleen (green) or intestine (blue) with age, regardless of the size of the change. Mutations that significantly changed with age, by beta binomial testing, are depicted by a star (spleen) or hashtag (intestine) on top of the bar. Group differences means (± SEM) were analyzed by one-way ANOVA with Tukey’s test (unless otherwise stated above). *p < 0.0332, **p<0.0021, ***p<0.0002,****p<0.0001.

### Tissue-specific selective pressures determine the fate of mtDNA mutations in *Polg*^D257A→WT^ mice

To determine why some tissues permit unconstrained expansion of early-life mtDNA mutations, while others do not, we examined the fate of mtDNA mutations with age. To do so, we used ultra-deep duplex sequencing to create a detailed mutational profile of 2- and 16-month-old *Polg*^D257A-flox^ and *Polg*^D257A→WT^ mice across five tissues, sequencing approximately 8.7 trillion raw bases, yielding over 75 billion duplex bases and detecting more than 100 million mutations, with an average depth of 100,000x (**Fig. 5F**). *Polg*^D257A→WT^ mice are uniquely suited for these analyses because they do not generate new mutations after recombination. Consequently, age-dependent changes in mutation frequency directly reflect the forces that govern the fate of pre-existing mutations, such as genetic drift and selection, rather than continued mutagenesis. We then compared the mutational profiles at the two time points to each other to determine which mutations tend to expand or contract over time, and elucidate their fate in a tissue-dependent manner. We classified these mutations as high, moderate, or low impact variants as determined by the Ensembl Variant Effect Predictor (VEP, *19*) and calculated how each impact class changed over time. To quantify these differences, we estimated the strength of purifying selection (s_log_2_FC) from the difference in median log_2_ fold change between synonymous and non-synonymous variants. Purifying selection was strongest in the spleen (0.365, 95% CI 0.300-0.409), followed by the heart (0.199, 95% CI 0.168-0.231), and intestine (0.077, 95% CI 0.057-0.099), whereas skeletal muscle (0.006, 95% CI −0.014 to 0.022) and liver (−0.039, 95% CI −0.068 to −0.0091) showed little evidence of selection (**Fig. 5G**). Importantly, the trajectory of this selection broadly matched the abundance of NBTx-positive cells in tissue and their physiological decline. For example, the spleen, which was devoid of NBTx-positive cells, selectively depleted mutations in the high impact category, suggesting that they undergo purifying selection; while the intestine, with many NBTx-positive cells, showed far less evidence of such discrimination, suggesting that the fate of mutations may be primarily guided by drift (**Fig. 5H**). These results indicate that tissues differ markedly in their capacity to engage in purifying selection against deleterious mtDNA variants, a property that is reflected in their physiology and clonal expansion.

### Selection is universal at critical regulatory elements

Although liver and intestine displayed little evidence of genome-wide purifying selection, mutations within three core regulatory elements showed strikingly stronger age-dependent depletion than mutations elsewhere in the genome (**Fig. S9A**). These elements: the displacement loop (D-loop), origin of light-strand replication (OL), and mTERF1 transcription termination site, govern mtDNA replication(*20*) and transcription(*21*), suggesting that mutations that disrupt these processes are subject to strong functional constraint irrespective of tissue identity (**Fig. 5I**). Consistent with this, in the intestine, the median log_2_ fold change at these positions was −0.51, while it was +0.12 in the rest of the genome (Mann-Whitney, p=2.2×10^−82^). Of the 85 genomic positions showing a significant, absolute >3-fold age-dependent change in mutation frequency in the intestine, 72 fell within these three regulatory regions, which is a strong enrichment over the genome-wide expectation (Fisher’s exact test, odds-ratio=92, p=4.7×10^−77^). The direction of change was overwhelmingly negative at these positions, with 62 of the 72 decreasing in frequency with age (binomial test, p=1.34×10^−10^), whereas the 13 positions elsewhere in the genome were evenly split and trended in the opposite direction (8 increased and 5 decreased). This contrast in directionality between regulatory and non-regulatory sites was itself significant (Fisher’s exact test, p=6.25x 10^−4^). The spleen displayed similarly strong selection at these sites but in addition, extended purifying selection throughout protein-coding genes, rRNAs and tRNAs. These observations underscore the importance of halting mutagenesis at 2 months of age, as unrecombined *Polg*^D257A-flox^ mice demonstrate a near universal increase in mutation frequency, obscuring the selective pressures exposed in *Polg*^D257A→WT^ mice (**Fig. S9B**). Taken together, these observations are consistent with a hierarchical organization of mtDNA quality control. Every tissue preserves the core machinery required for mitochondrial genome maintenance, whereas only a subset extends purifying selection across the remainder of the mitochondrial genome. For example, despite relatively weak selection across the surrounding sequence, mutations within the core mTERF1 recognition motif(*21*) that terminates transcription were consistently depleted with age, highlighting the importance of preserving this essential regulatory element (**Fig. 5J**). Interestingly, we also noticed that the nucleotide corresponding to the pathogenic human MELAS mutation m.3243A>G(*22*) exhibited the weakest depletion within the core motif, raising the possibility that reduced selective pressure at this position contributes to the prevalence of this mutation in human disease.

### Mitochondrial fusion controls the fate of somatic mutations in mammalian cells

The striking differences in purifying selection between tissues raised the question of which cellular mechanisms determine their ability to eliminate deleterious mtDNA mutations. One candidate is mitochondrial fusion. Previously, inhibition of fusion was shown to promote purifying selection against high-heteroplasmy variants in the Drosophila germline(*23*) and muscle(*24*) by exposing dysfunctional mitochondria to mitophagy. We therefore asked whether inhibiting mitochondrial fusion could also promote the selective removal of rare early-life mutations, which are initially innocuous, but could drive aging later in life through their expansion. To test this hypothesis, we isolated primary fibroblasts from WT, *Mfn1*^loxp^, *Polg*^D257A-flox^, and *Polg*^D257A-flox^; *Mfn1*^loxp^ mice. We then recombined the *Polg* allele in these cells to stop mutagenesis (**Fig. S10A,B**), deleted *Mfn1* to inhibit mitochondrial fusion (**Fig. 6A-C, Fig. S10C**), and monitored the fate of mtDNA mutations over one month in cell culture. While high-impact variants readily accumulated in *Polg*^D257A→WT^ control fibroblasts (p=0.03), they were selectively depleted following *Mfn1* deletion (p=0.002, **Fig. 6D**). This effect was particularly pronounced for protein-truncating mutations, whereas D-loop variants were unaffected (**Fig. 6E**), indicating that loss of fusion selectively removes mutations that compromise oxidative phosphorylation rather than indiscriminately eliminating mtDNA genomes. We next asked whether this mechanism extends to pathogenic human mtDNA mutations. Consistent with this idea, CRISPR-Cas9 based deletion of *MFN1* in human osteosarcoma cells carrying the mitochondrial common deletion (∼90% heteroplasmy) reduced deletion burden by approximately 85% over three months, while treatment with scrambled gRNAs did not (**Fig. 6F**). Finally, we tested whether inhibiting fusion could confer selective capacity on a tissue that normally lacks it. Following simultaneous recombination of the *Polg*^D257A-flox^ allele and deletion of *Mfn1 in vivo*, NBTx-positive cells in the intestine fell from 16% in *Polg*^D257A→WT^ mice to only 1.3% in *Polg*^D257A→WT^; *Mfn1*^−/−^ mice (p=0.0089, **Fig. 6G, Fig. S10D**), suggesting that mitochondrial fusion is a key determinant of mtDNA quality control. Most importantly though, they demonstrate that even though pathogenic mtDNA mutations may be seeded early in life, the ultimate fate of these mutations is not fixed. Instead, it can be reshaped by manipulating the cellular mechanisms that determine whether deleterious genomes persist or are eliminated.

**Figure 6.**
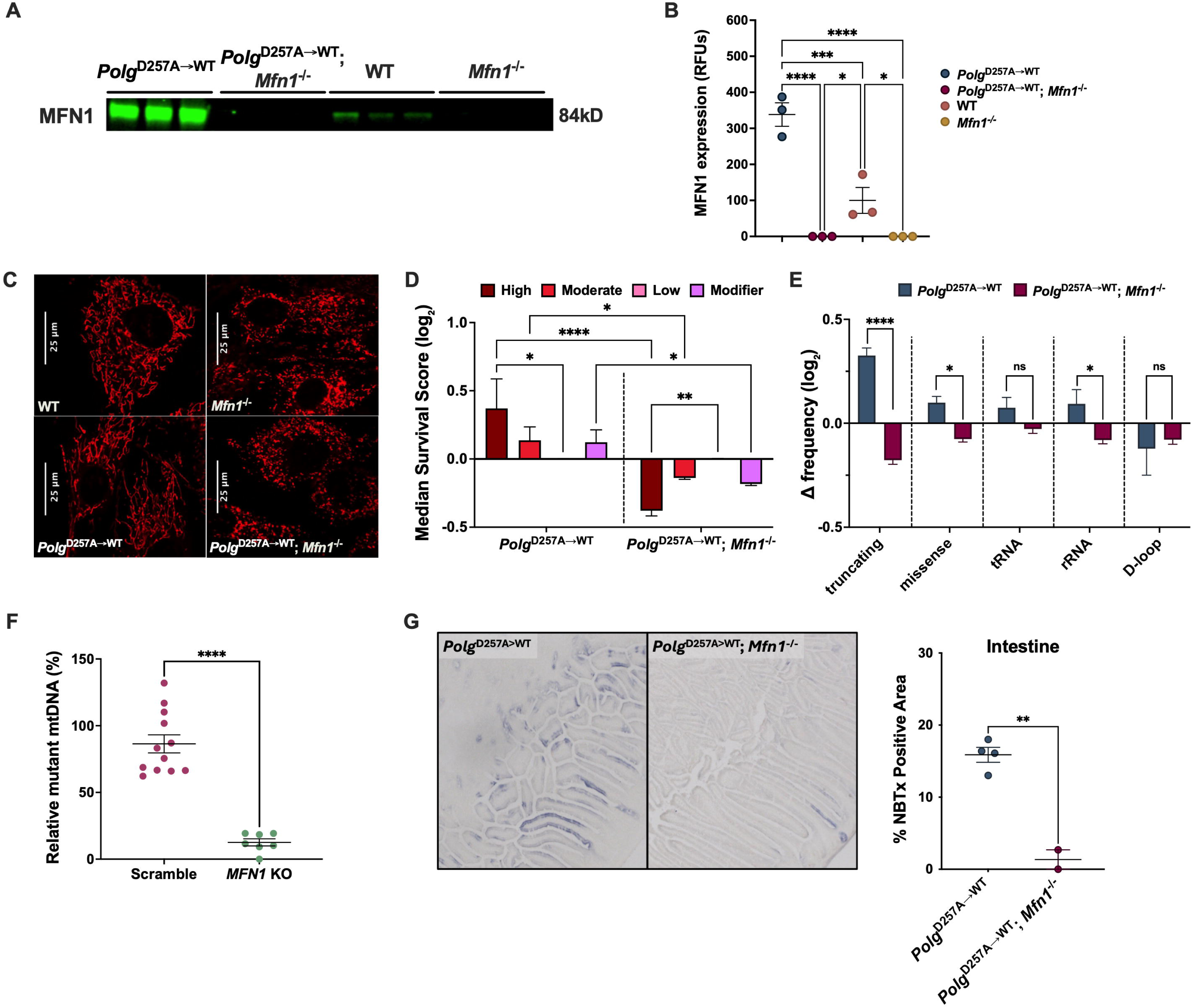
Inhibition of mitochondrial fusion drives purifying selection against deleterious mtDNA mutations in somatic cells and tissues. (**A,B**) Representative Western blot (**A**) and quantification (**B**) of MFN1 protein in fibroblasts derived from *Polg*^D257A→WT^, *Polg*^D257A→WT^; *Mfn1*^−/-^, WT, and *Mfn1*^−/-^ mice, confirming loss of MFN1 upon recombination and elevated MFN1 in *Polg*^D257A→WT^ cells relative to WT (n=3/group). (**C**) Confocal images at 63x magnification of mitochondrial morphology in WT, *Mfn1*^−/-^, *Polg*^D257A→WT^, and *Polg*^D257A→WT^; *Mfn1*^−/-^ fibroblasts, showing fragmentation of mitochondria upon loss of *Mfn1*. Scale bars, 25 μm. (**D,E**) Selection against mtDNA variants in *Polg*^D257A→WT^ and *Polg*^D257A→WT^; *Mfn1*^−/-^ fibroblasts passaged for one month, computed from duplex sequencing as the median log_2_FC in variant frequency relative to synonymous variants. (D) Variants stratified by predicted functional impact. (**E**) Variants stratified by functional class, showing that loss of *Mfn1* selectively removes protein-truncating variants, whereas depletion of D-Loop variants is comparable between genotypes. (**F**) Ratio of the common deletion to total mtDNA (deletion/12S) in human cybrid cells three months after *MFN1* knockout compared to scramble control, measured by qPCR. *MFN1* loss reduces the deletion burden by 85% (n=10-12 clones/group). (**G**) Representative NBTx staining and quantification of intestinal tissue from 8-month-old *Polg*^D257A→WT^ and *Polg*^D257A→WT^; *Mfn1*^−/-^ mice, showing near-complete loss of clonally expanded, energy-deficient cells upon *Mfn1* deletion (n=2-4/group). Group differences means (± SEM) were analyzed by one-way ANOVA with Tukey’s test (**B**), by two-way ANOVA with Fisher’s LSD test (**D,E**), or by Welch’s t-test (**F,G**). *p<0.0332, **p<0.0021, ***p<0.0002, ****p<0.0001.

## DISCUSSION

A central question in the biology of aging is when the cellular and molecular events that ultimately drive physiological decline first arise. By restricting mitochondrial mutagenesis to the first two months of life, we demonstrate that early-life mtDNA mutations are sufficient to drive a wide range of age-related pathologies that often do not become apparent until a year later. These observations establish that mtDNA mutations can remain functionally silent for prolonged periods before contributing to disease, consistent with the idea that they act as “temporal seeds” whose pathogenic potential is only realized after years of clonal expansion. Thus, in some tissues the biological trajectory toward aging may be established almost as soon as development is complete.

An important caveat of our study though, is that it only establishes sufficiency, not necessity. Although early-life mtDNA mutations clearly have the capacity to drive age-related pathology, our experiments do not demonstrate that this is the principal mechanism underlying physiological aging. Definitive proof for that idea will require experimental strategies that selectively eliminate early-life mtDNA mutations while permitting later-life mutagenesis to proceed normally. Our observations, suggest that temporary inhibition of mitochondrial fusion may be one way to do that, while other models of mitochondrial mutagenesis may be useful as well(*25*). Nevertheless, several arguments suggest that a similar process may occur in WT animals. First, because multiple tissues show a lack of selection, mutations that are acquired early in life have the greatest opportunity to undergo clonal expansion during subsequent decades, making them prime candidates to dominate aged tissues. Second, previous research has shown that the mtDNA mutation spectrum found in aged human intestinal crypts closely resembles the spectrum generated early in life(*9*), an observation that is remarkably consistent with our work.

We also identified several tissues that are inherently resilient to early life mutations. These tissues seem to defy the idea that mutations are irreversible, permanent additions to our genome. By exploiting the malleability of the population of mtDNA molecules inside cells, they selectively remove genomes that have the potential to cause cellular aging later in life. These observations echo findings in flies, where a high heteroplasmy mutation present throughout the organism was purged from the germline(*23*) and muscle(*24*) tissue by manipulating mitochondrial fusion. Our data builds on these findings by demonstrating that mammals have similar capabilities, and reveal its full potential in the context of aging: as a mechanism to rejuvenate cells at a genetic level.

These findings support the idea of mitochondrial fusion as a double-edged sword. On the one hand, fusion promotes mitochondrial function by allowing content from WT mitochondria to complement mutant mitochondria, while on the other hand, this complementation process prevents respiratory defects that can trigger mitochondrial quality control. Thus, mitochondrial dynamics may provide a mechanism through which cells balance the immediate benefits of functional complementation against the long-term benefits of genetic quality control (**Fig. S11**). Different tissues may exploit this balance differently. For example, proliferating *Polg*^D257A→WT^ fibroblasts significantly increase Mfn1 expression, favoring complementation (**Fig. 6A,B**), whereas the highly ordered and spatially constrained mitochondrial populations of tissues such as the heart(*26*) may facilitate segregation and removal of dysfunctional mitochondria. Consistent with this model, inhibition of fusion in our experiments prevented clonal expansion in the intestine of recombined mice and selectively eliminated deleterious mtDNA mutations in *Polg*^D257A→WT^ fibroblasts. Notably, this selection was not indiscriminate. Loss of fusion depleted protein-truncating variants, which disrupt oxidative phosphorylation, while leaving D-loop variants unchanged (**Fig. 6E**), implicating a mechanism that senses respiratory dysfunction, most likely mitophagy.

Our observations also have important therapeutic implications. Because of their reputation as permanent age-related changes, most approaches to combat mtDNA mutations have traditionally focused on reducing the formation of new mutations(*27, 28*). However, if pathogenic mtDNA mutations are acquired early in life, treatments to prevent mutations are unlikely to help the middle-aged and elderly, and may only prove effective in young individuals. Our findings suggest a fundamentally different strategy: rather than preventing mutations from arising, it may be possible to selectively eliminate pathogenic mtDNA genomes after they have already formed. Our experiments identify mitochondrial fusion as one mechanism through which this can be achieved. However, a deeper understanding of the molecular pathways that mediate this selective quality control, together with identifying the classes of mtDNA mutations that are preferentially removed, may reveal entirely new therapeutic strategies capable of slowing, preventing, or perhaps even reversing age-related mitochondrial dysfunction.

## Acknowledgements

Mass spectrometry experiments for this project was done on instruments in the USC Mann Multi-Omics Mass Spectrometry Core. Sectioning for H&E was done with the help of the Translational Pathology Core Laboratory at UCLA. Graphical illustrations were created using BioRender. Circle plot was generated using the Circa software.

## Author Contributions

Conceptualization: MV (mouse model and study design), SJS (study design)

Methodology: MV, SJS

Investigation: SJS, MV, EH, LC, CSC, SL, HA, BMV, JW, IV, MAT, WC, and ARV

Visualization: SJS, MV

Data Curation: JFG

Software: JFG

Formal Analysis: JFG, BAB, SJS, and MV

Supervision: MV, SJS

Funding acquisition: MV

Writing – original draft: MV, SJS

Writing – review & editing: BMV, SRK, ARV, JFG, WC, BAB, PC, MAT, JW, SJS, and MV

## Materials and Correspondence

Correspondence and requests for materials should be addressed to.

## Data, code and materials availability

All data needed to evaluate and reproduce the conclusions in the paper are present in the paper and/or the Supplementary Files. A modified version of the standard Duplex-Seq pipeline was used to analyze the duplex sequencing data and has been archived on Zenodo at https://doi.org/10.5281/zenodo.22943789. Raw data are in the process of being deposited to SRA. All other code used to analyze data for this manuscript can be found at https://github.com/sshemtov1/temporal_control_mito_mut.

## Materials & Methods

### Generation of *Polg*^D257A-flox^ mice

To generate mice carrying a conditional proofreading-deficient allele of *Polg*, targeted embryonic stem (ES) cell clones were produced by homologous recombination at the endogenous *Polg* locus. Correctly targeted ES cell clones were injected into C57BL/6 albino blastocysts, which were subsequently implanted into pseudopregnant CD-1 females. Chimeric founder animals were identified based on coat color contribution and bred to C57BL/6 females to assess germline transmission. Germline transmission of the targeted allele was confirmed by genotyping of offspring. Independent founder lines were established from both ES cell clones, yielding *Polg*^D257A-flox/+^ mice. Founder lines were generated by Cyagen Biosciences (TKX-190523). Male *Polg*^D257A-flox/+^ mice were then crossed with female WT C57Bl6/J mice (JAX #:000664) to expand the colony. Male and female heterozygous *Polg*^D257A-flox/+^ mice were later crossed to generate homozygous *Polg*^D257A-flox^ mice that were initially used to characterize the phenotype of our mutant mouse model and determine whether the mutant transgene resulted in a premature aging phenotype similar to that observed in unconditional *Polg*^D257A^ mice. At 21 days, mice were weaned, ear-punched and one PCR reaction was run to genotype the *Polg* allele of each mouse using primers: F: 5’-GTGTGCATACATTCTCATCTTGGC-3’ and R: 5’-ACGAATAGGACCAACCCCTTGTGTC-3’. PCR reactions were run on agarose gels by gel electrophoresis and imaged before analysis.

### Generation of *Polg*^D257A-flox^ x Cre^ERT2^ Mice

Initially, we crossed male *Polg*^D257A-flox/+^ mice to female ROSA26-Cre-ER^T2^ mice(*29*) (JAX #: 008463) and later crossed mice heterozygous for both *Polg*^D257A-flox^ and ROSA26-Cre-ER^T2^ to generate mice heterozygous or homozygous for either or both alleles. The ROSA26-Cre-ER^T2^ locus was genotyped using two separate reactions, one for the presence of the WT allele (primers: F: 5’-CTGGCTTCTGAGGACCG-3’ and R: 5’-CCGAAAATCTGTGGGAAGTC-3’) and one for the presence of the transgene (primers: F: 5’-CGTGATCTGCAACTCCAGTC-3’ and R: 5’-AGGCAAATTTTGGTGTACGG-3’). Once we determined that the ROSA26-Cre-ER^T2^ model was not providing us with sufficient recombination of the *Polg*^D257A-flox^ allele across all tissues (see strategies tested below), we then crossed male *Polg*^D257A-flox/+^ mice to female UBC-Cre-ER^T2^ mice(*30*) (JAX #: 007001), which were maintained as hemizygotes. We then crossed *Polg*^D257A-flox/+^ and *Polg*^D257A-flox/+^; UBC-Cre-ER^T2/+^ to generate mice homozygous for the *Polg*^D257A-flox^ allele. The UBC-Cre-ER^T2^ locus was genotyped in a single reaction using four primers, one set for the presence of the WT allele (primers: F: 5’-CTAGGCCACAGAATTGAAAGATCT-3’ and R: 5’-GTAGGTGGAAATTCTAGCATCATCC-3’) and one set for the presence of the transgene (primers: F: 5’-GACGTCACCCGTTCTGTTG-3’ and R: 5’-AGGCAAATTTTGGTGTACGG-3’).

### Cre Recombination Strategies

When mice reached 2-months of age, we began administering tamoxifen through various strategies in order to find one that resulted in 100% recombination in all tissues. Mice were euthanized two weeks after tamoxifen administration concluded to determine efficiency of recombination. Recombination of all tissues was assessed by RNA extraction (Ribopure Purification Kit, Invitrogen #AM1924), cDNA synthesis (SuperScript III First-Strand Synthesis System, Invitrogen #18080051), and PCR using primers: F: 5’-TGCTTGGCAGAGGGAACCT-3’ and R: 5’-GTATCGAGAAAACGCATCCG-3’, before being sent out for Sanger Sequencing and the traces subsequently analyzed. Tamoxifen (MilliporeSigma, cat# T5648) was resuspended in corn oil at a concentration of 20mg/mL, 100 mg of (Z)-4-Hydroxytamoxifen (4-OH tamoxifen; MilliporeSigma, cat# H7904) was first resuspended in 1mL of 100% ethanol before being added to 9mL of corn oil, for a final concentration of 10mg/mL. Tamoxifen solutions were administered by oral gavage or by intraperitoneal injection daily for 5 days at 50mg/kg (4-OH tamoxifen), 100mg/kg (tamoxifen and 4-OH tamoxifen), or 200mg/kg (tamoxifen). Custom tamoxifen chows were purchased at 250mg tamoxifen/kg diet and 500mg tamoxifen/kg diet from Envigo and provided to the mice *ad libitum* for 4 weeks. Ultimately, 200mg/kg tamoxifen by oral gavage was chosen for the remainder of the study. While heterozygous *Polg*^D257A-flox^; UBC-Cre^ERT2/+^ mice were 100% recombined with a single week treatment regimen, we found that homozygous *Polg*^D257A-flox^; UBC-Cre^ERT2/+^ mice required an extra week of treatment. Therefore, experimental animals were gavaged daily for 5 days, followed by one week of recovery, and then another 5-day treatment.

### Experimental Animals

In order to prevent mtDNA mutations from being inherited, male *Polg*^D257A-flox/+^ mice were continuously bred with female UBC-Cre^ERT2/+^ mice in order to prevent females from passing down mtDNA genomes with a significant mutant load to their offspring. To generate homozygous *Polg*^D257A-flox^ mice, *Polg*^D257A-flox/+^ females used in these crosses always were derived from a male *Polg*^D257A-flox/+^ x female UBC-Cre^ERT2/+^ cross to limit this confounder. Mice were assayed and euthanized at 2-months or 16-months of age for experimental studies, except for *in vivo Mfn1* knockout experiments, for which mice were euthanized at 8-months of age. Conditional *Mfn1*^loxp^ mice were purchased from JAX (#: 026401)(*31*) and crossed into *Polg*^D257A-flox^ ; UBC-Cre^ERT2^ backgrounds. Mice used in *in vivo Mfn1* knockout experiments were homozygous for *Polg*^D257A-flox^ and *Mfn1*^loxp^, and heterozygous for UBC-Cre^ERT2^. The *Mfn1* allele was genotyped using primers: F: 5’-TGGTAATCTTTAGCGGTGCTC-3’ and R: 5’-GGAGGACTTTATCCCACAGC-3’). Body composition was measured prior to euthanasia using the Bruker LF90 II Minispec (Bruker Optics). Mice were placed and loosely restrained within an acrylic cylinder. The cylinder was placed inside instrument and fat, lean, and fluid mass was measured. Animals were returned to their original cages as soon as the scan was completed. Both male and female mice were evaluated for organismal and physiological phenotypes. Physiological outcomes were not significantly different between males and females, so the data were plotted together. If appropriate, data are shown separately by sex (lean mass). Because these phenotypes were concordant between sexes, and to limit animal numbers in accordance with 3R principles, molecular profiling and staining were performed in male mice. All animal research was approved by USC’s Institutional Animal Care and Use Committee (protocol #20930). Animals were scored by our lab and USC’s Department of Animal Resources veterinarians to establish humane endpoints. Mice were euthanized by CO_2_ exposure.

### Muscle Function

Grip strength was assessed using a horizontally mounted bar attached to a sensor (TSE-Systems, 303500-M/E1) designed to measure limb strength. Each mouse was allowed to securely grasp the bar before being swiftly and steadily pulled backward. Measurements were recorded only if the mouse let go with both forelimbs at the same time. Ten measurements were taken per animal and averaged to obtain a final value. For endurance measurements, mice were acclimated to the treadmill instrument (TSE 303401-M-04/C) 24 hours before testing. During acclimation, animals were placed on the stationary treadmill for five minutes, followed by an increase in speed to 2m/min for five additional minutes. Mice were then tested once each day on the two days following acclimation. The treadmill protocol consisted of 1 m/min for 1 minute, followed by a 1m/minute increase every minute until exhaustion. Time of exhaustion was recorded in seconds. Mice that refused to run received a mild electrical stimulus at the back of the treadmill. The values recorded for each mouse over both test days was averaged and reported.

### Echocardiography

One week prior to dissection, mice were transported to the molecular imaging center at USC for echocardiography using the VisualSonics Vevo 3100, MX 550 transducer 22-55 MHz. Mice were anesthetized with 2% isoflurane and a depilatory cream (Nair) was used to remove fur prior to echocardiography. Mice were placed in the supine position onto the warmed platform to maintain optimal physiological conditions and their limbs were taped onto the metal EKG leads. Heart rates were monitored and generally maintained at 400–500 beats per minute. Warmed echocardiography gel was placed on the shaved chest and the heart was imaged with a 30 MHz transducer. By placing the transducer along the long-axis of LV and directing it to the right side of the neck of the mouse, two-dimensional LV long-axis can be obtained. The transducer was then rotated clockwise by 90°, and the LV short-axis view was obtained. Transmitral inflow Doppler spectra were recorded in an apical 4-chamber view by placing the sample volume at the tip of the mitral valves. After the scans were concluded, the residual gel was removed, and the mouse was returned to the cage for recovery. Images were subsequently analyzed using the VevoLab software.

### μCT Scanning

One week prior to dissection, mice were transported to the molecular imaging center at USC for μCT scan using the Rigaku CT Lab GX, with the following settings: 90kV, 88uA, High Resolution at 45mm FOV, Voxel Size: 90μm. Mice were anesthetized with 2% isoflurane and placed in the prone position on the imaging platform. Mice were taped loosely onto the imaging platform to minimize motion artifact caused by animal movement (breathing or heartbeat) during data acquisition. The scans lasted 4-5min, and once complete, animals were returned to their original cages for recovery.

### Tissue Collection and Analysis

Upon euthanasia, mice were measured, photographed, and the heart was punctured for blood draws for blood composition experiments and plasma isolation. Tissues were dissected, placed on a grid sheet for measurements and photography, and cut into various pieces. One piece was frozen in Optimal Cutting Temperature (OCT) compound, and three pieces were flash frozen in liquid nitrogen for DNA, RNA, and protein extractions. Testes, spleen, and skin were fixed in formalin at room temperature overnight before being washed twice with PBS and stored at 4°C in 70% EtOH. Tissues were then embedded in paraffin and sectioned (4μm) before H&E staining, performed by the Translational Pathology Core Laboratory at UCLA. Sections were imaged using the ECHO Revolve Microscope. Blood from cardiac punctures was immediately collected in EDTA tubes for plasma isolation (15-minute spin at 2,000g at 4°C). Plasma was stored at −80°C until experimentation. Pro-inflammatory cytokines were measured in the plasma with a commercial immunoassay, V-PLEX Proinflammatory Panel 1 (mouse) Kit (Meso Scale Discovery, K15048D-1) and plasma GDF15 was measured by ELISA (Proteintech Group, Inc. cat #: KE10082).

### DNA extraction & Duplex-Sequencing

Tissue and cell line DNA was extracted using the Qiagen DNeasy Blood and Tissue Kit (cat# 69506), quantified by Qubit 1x dsDNA hs assay kit, and stored at −20°C. 500ng of DNA per sample was fragmented using the Biorupter Pico (30 seconds ON, 90 seconds OFF, 7 cycles) and size was measured using the Agilent 4200 TapeStation instrument (fragment size ∼200bp). Published protocols(*13, 32, 33*) were followed with minor adjustments. Briefly, DNA was end-repaired and adapters were ligated to DNA fragments using the Ultra II DNA Library Preparation Kit from NEB (cat# E7103) samples were cleaned up and libraries were quantified by qPCR using SYBR Green iTaq Supermix (Bio-Rad cat#: 1725124) compared to previously generated libraries with target family sizes and on-target efficiencies. This quantification enables us to input the correct number of molecules into the pre-enrichment PCR depending on mtDNA content for each sample to sequence at the target parameters (depth, target raw reads, family size, on-target reads). Libraries were then enriched for mtDNA sequences using the xGen Hybridization Capture Assay (IDT) with a custom Discovery Pool of biotinylated probes (**Suppl. File 1**), along with their protocol. Final libraries were quantified, pooled and sequenced on a NovaSeq X Plus 25B. A modified version of the standard Duplex-Seq pipeline was used to analyze the sequencing data (Zenodo https://doi.org/10.5281/zenodo.22943789). Raw data are being deposited to SRA.

### Mutation analysis and variant functional impact classification

#### Tissues

Variant call format (VCF) files generated from the duplex sequencing pipeline were analyzed using R (version 4.6.1). Mutation files from individual biological replicates were first combined by age group (2-month-old *Polg*^D257A-flox^ mice and 16-month-old *Polg*^D257A→WT^ mice). Mutations were matched across replicates based on chromosome, genomic position, reference allele, alternate allele, and variant type. Replicate-specific mutation frequencies were retained for each mutation, and average mutation frequency and sequencing coverage were calculated across replicates. To reduce inter-animal variability, mutations exhibiting high variability between replicates were filtered out using a mutation frequency range threshold of 0.005. Filtered datasets from the two age groups were then merged, and age-associated changes in mutation frequency were quantified by calculating ratios and log_2_ fold changes (log_2_FCs) between 16-month and 2-month samples. Final processed datasets were exported and annotated using the Variant Effect Predictor (VEP; Ensembl) to assign predicted functional impact. For each tissue and impact class, the median log_2_FC was computed across variants. Uncertainty in the median was estimated using non-parametric bootstrap resampling. Log_2_FC values within each tissue and impact group were resampled with replacement 2,000 times, and the median was recalculated for each bootstrap replicate. The 2.5th and 97.5th percentiles of the bootstrap distribution were taken as the 95% confidence interval.

#### Cell Lines

VCF files generated from the duplex-sequencing pipeline were analyzed in R 4.6.1. Mutation files from individual biological replicates were annotated using VEP to assign predicted functional impact, and variants receiving multiple transcript annotations were collapsed to unique loci, retaining the most severe predicted impact. We analyzed only variants that were reliably present at baseline (baseline mutant-allele frequency >4×10^−5^), so that changes over time reflected genuine gains or losses rather than detection noise. Each replicate cell line and timepoint was processed separately, and all statistics treated the cell line as the unit of replication (three lines per genotype), so error bars and significance reflect variability between independent cell lines rather than between individual variants. For each variant, time- or recombination-associated change in frequency was summarized as a unified survival score that incorporates both partial reduction and complete loss. Given the high duplex sequencing depth (median coverage ∼1×10^5^), a variant detected at baseline but absent at the later timepoint was treated as a genuine complete loss rather than a detection failure. Variants still detected at the later timepoint were scored as the log_2_FC in frequency (log_2_(freq_late / freq_pre)), while completely lost variants, for which this ratio is undefined, were left-censored at the assay detection limit and scored as log_2_[(0.5 / C) / freq_pre], where C is the replicate’s median later-timepoint coverage. This places frequency reduction and complete elimination on a common scale and avoids the survivorship bias that arises when lost variants are discarded. To remove the confounds of cell culture that result in a global downward shift in frequency introduced by artificial bottlenecks, unified scores were referenced to synonymous variants as an empirical neutral baseline. For each replicate and timepoint, we computed the median synonymous score and its median absolute deviation (MAD, unscaled), and summarized each impact class by the number of variants, the complete-loss rate, and the median and mean unified score, together with a synonymous-referenced survival difference (median -synonymous median) and a MAD-normalized selection score, (median - synonymous median) / synonymous MAD. Because the synonymous baseline differs substantially between conditions and timepoints, cross-condition and cross-replicate comparisons used the synonymous-referenced values rather than the raw scores. Negative scores indicate depletion relative to the neutral baseline (purifying selection).

#### Estimation of selection strength – Tissues

For each tissue, log_2_FCs between 16-month and 2-month animals (as above) and selection strength was determined by comparing synonymous and non-synonymous variants (as predicted by VEP). Synonymous variants were treated as an empirical neutral reference, while non-synonymous variants included missense, nonsense, frameshift, and in-frame insertion or deletion mutations. A trajectory-based coefficient was defined as the difference between the median log_2_FC of synonymous and non-synonymous variants:

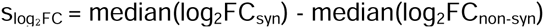

Positive values of slog_₂_FC indicate purifying selection acting against non-synonymous variants. This metric represents a relative, trajectory-based measure of selection strength, with synonymous variants serving as an empirical neutral reference rather than implying a direct estimate of population-genetic selection coefficients. Uncertainty in selection coefficient estimates was estimated using non-parametric bootstrap resampling. For each tissue, synonymous and nonsynonymous log_2_FC values were resampled with replacement 2,000 times, and the selection coefficient was recalculated for each bootstrap replicate as the difference of medians. The 2.5th and 97.5th percentiles of the bootstrap distribution were taken as the 95% confidence interval. This approach avoids distributional assumptions and is robust to skewed distributions and unequal variances between functional classes.

#### Functional-class selection analysis – Cell Lines

To test whether selection targets specific functional categories, variants were assigned to classes by VEP biotype and consequence: protein-truncating (stop-gain, frameshift, stop-loss, or start-loss), missense, tRNA (mt_tRNA biotype), rRNA (mt_rRNA biotype), and D-Loop (control-region/intergenic), with synonymous variants as the neutral reference. Because protein-disrupting variants tend to arise at lower baseline frequency, and lower-frequency variants are more readily lost irrespective of impact, functional-class effects were estimated with baseline frequency as a covariate. For each replicate, the unified survival score was modeled by ordinary least squares as a function of functional class and log_10_ baseline frequency, with synonymous variants as the reference category (u ∼ functional class + log_10_[baseline frequency]). Each functional class coefficient therefore represents the difference in unified survival relative to synonymous variants at the same modeled baseline frequency. Negative coefficients indicate relative depletion, consistent with selection against that class. For each genotype, coefficients were summarized across the three replicate lines using the mean and SEM.

### Position-level beta-binomial testing of age-associated mutation frequencies

Mutation calls were collapsed to one observation per genomic position by summing mutation counts at a given position. Per-animal coverage was taken once and positions absent from the input were not treated as zero-mutation observations. At each position, mutation counts and total coverage were analyzed with a beta-binomial model to accommodate overdispersion among biological replicates. A likelihood-ratio test compared a null model with one shared mutation frequency across age groups with an alternative model allowing separate 2-month and 16-month frequencies while retaining a shared position-specific dispersion parameter. P values were obtained from a chi-square distribution with 1 degree of freedom and adjusted across tested positions using the Benjamini-Hochberg procedure. We subsequently calculated the average mutation frequency for each age group, the 16-month/2-month fold change and fold-change threshold (absolute fold change of 2 or 3).

### Untargeted Proteomics

#### Sample Processing and Liquid Chromatography-Tandem Mass Spectrometry

Proteomic analysis was conducted at the USC Alfred E. Mann Multi-Omics Mass Spectrometry Core (Director: Whitaker Cohn, Ph.D). The tissues were homogenized in a lysis buffer containing 12mM sodium lauroyl sarcosinate, 0.5% sodium deoxycholate, 50mM triethylammonium bicarbonate (TEAB), and Halt™ Protease and Phosphatase Inhibitor Cocktail (Thermo Fisher Scientific) via bead beater at 4°C. Protein concentration was determined using a BCA Protein Assay. An aliquot of each sample (10µg) was reduced and alkylated using 10mM tris(2-carboxyethyl)phosphine and 40mM chloroacetamide for 20 minutes at 95°C. For buffer exchange, SP3 SpeedBead Magnetic Carboxylate-Modified Particles (250µg total, Cytiva; Cat# 65152105050250 and 4515210505250, mixed 1:1) were added. Ethanol was added to a final concentration of 55%, and samples were gently agitated for 15 minutes to facilitate protein binding, followed by three washes with 80% ethanol. Proteins were eluted from the beads with 50µL of 50mM TEAB containing 0.5µg of mass spectrometry-grade Trypsin/Lys-C Mix (Promega, Cat# V50711) and digested overnight at 37°C for 16 hours. Peptide aliquots (200ng) were injected onto a µPAC Neo Plus C18 column (50cm; Thermo Fisher Scientific, Cat# COL-NANO050NEOB) equilibrated in solvent A (0.1% formic acid) and separated using a Vanquish Neo UPLC system (Thermo Fisher Scientific) with an optimized 49-minute gradient of solvent B (acetonitrile/water/formic acid, 80/20/0.1, v/v/v) as follows (min/%B/flow rate µL/min): 0/4/0.75, 0.1/5/0.75, 4.1/7.2/0.3, 27.6/20/0.3, 41.8/35/0.3, 42.8/55/0.3, 43/99/0.3, 47/99/0.3, 47.2/99/0.75, 49/99/0.75. Eluted peptides were introduced into a nanospray ionization source coupled to an Orbitrap Astral mass spectrometer (Thermo Fisher Scientific) operated in data-independent acquisition mode. Full MS scans were acquired in the Orbitrap (m/z 380–980; AGC target 500%; maximum injection time 3 ms; resolution 240,000 at m/z 200). MS2 scans were acquired in the Astral analyzer using 2 m/z isolation windows, 25% HCD collision energy, AGC target 300%, maximum injection time 3 ms, and a resolution of 17,500 at m/z 200. Raw proteomic data were searched against a custom database encompassing the complete mouse proteome and common mass spectrometry contaminants using CHIMERYS in Proteome Discoverer (Version 3.3, Thermo Fisher Scientific). Static modifications included carbamidomethyl (+57.021) on cysteine (C) and dynamic modifications included oxidation (+15.995) on methionine (M). Detection of b+ and y+ fragment ions from tryptic peptides were utilized to confidently identify and quantify peptides belonging to proteins within the provided database.

#### Proteomic Data Processing and Statistical Analysis

Raw protein abundance values were extracted from mass spectrometry-derived intensity columns. Proteins detected in fewer than four of six samples in all experimental groups were excluded. Remaining abundance values were log2-transformed prior to statistical analysis. Between-sample normalization was performed using quantile normalization (limma::normalizeBetweenArrays) to align global intensity distributions across samples. Differential protein abundance was assessed using linear models with empirical Bayes moderation implemented in the limma package. Group-specific design matrices were constructed without intercept terms, and contrasts were specified to compare *Polg*^D257A-flox^ vs WT, *Polg*^D257A→WT^ vs WT, and *Polg*^D257A-flox^ vs *Polg*^D257A→WT^ groups. Moderated t-statistics were calculated and p-values were adjusted using the Benjamini–Hochberg FDR procedure. Proteins with FDR<0.05 were considered differentially expressed. Nnt protein was excluded from the interpretation of the results, due to some mice carrying a heterozygous allele. Analysis was redone to exclude replicates heterozygous for Nnt, and no difference in significant proteins was observed, so we decided to include all replicates (n=6) and remove the protein from interpretation. Over-representation analysis (ORA) of Reactome pathways was performed on differentially expressed proteins using the ReactomePA package (enrichPathway). This analysis was carried out independently for each of the three contrasts in both heart and liver tissue. For each contrast, protein gene symbols were mapped to Entrez gene identifiers against the Mus musculus annotation database (org.Mm.eg.db) using clusterProfiler::bitr. To restrict inference to proteins that were actually measurable in the experiment, the background gene universe was defined as all quantified proteins retained after detection filtering and tested in the differential-abundance model for that contrast, rather than the entire mouse genome. ORA was performed separately on the full set of significant DEPs and on the upregulated and downregulated subsets separately, defined by the sign of the log_2_FC. Reactome pathways with a Benjamini–Hochberg p_adj_<0.05 were considered significantly enriched. Fold enrichment was calculated as the ratio of the proportion of DEPs annotated to a given pathway (GeneRatio) to the proportion of background proteins annotated to that pathway (BgRatio). All analyses were performed in R 4.6.1. Differential abundance was assessed with limma (v3.68.5), and Reactome over-representation analysis with clusterProfiler (v4.20.0) and ReactomePA (v1.56.0), using org.Mm.eg.db (v3.23.0) and reactome.db (v1.96.0) for annotation. Raw and analyzed data are available in **Suppl. File 2**.

### Untargeted Metabolomics

#### Sample Processing and Liquid Chromatography-Tandem Mass Spectrometry

Tissues were placed in 10μL ice-cold 4:1 LC-grade Methanol:Ultrapure-water (Fisher) per mg tissue and homogenized using a bead homogenizer (Bead Mill 4, Fisherbrand). Tissue homogenates were centrifuged at 4°C at 21,300g for 10 minutes and the metabolite-containing supernatants were collected. 100μL of supernatant were dried down per sample using a vacuum centrifuge and stored at −80°C until further processing. An aliquot of each sample (3μL) was injected onto an XBridge BEH Amide column (2.1×150 mm, 2.5μm particle size, 130Å pore size; Waters Corporation) and separated using a Vanquish UHPLC system (Thermo Fisher Scientific). Mobile phase A consisted of 95:5 water/acetonitrile containing 20mM ammonium acetate and 20 mM ammonium hydroxide (pH 9.45), and mobile phase B consisted of acetonitrile. Chromatographic separation was performed at a flow rate of 150μL/min using the following gradient: 0-2 min, 85% B; 3 min, 80% B; 5 min, 80% B; 6 min, 75% B; 7 min, 75% B; 8 min, 70% B; 9 min, 70% B; 10 min, 50% B; 12 min, 50% B; 13 min, 25% B; 16 min, 25% B; 18 min, 0% B; 23 min, 0% B; 24 min, 85% B; and 30 min, 85% B for column re-equilibration. The column effluent was introduced by electrospray ionization into a Q Exactive quadrupole-Orbitrap mass spectrometer (Thermo Fisher Scientific). Source parameters were as follows: spray voltage, +3.8kV in positive ion mode and −3.5kV in negative ion mode; capillary temperature, 300°C; auxiliary gas heater temperature, 360°C; sheath gas, 40 arbitrary units; auxiliary gas, 10 arbitrary units; sweep gas, 2 arbitrary units; and S-lens RF level, 50. Individual study samples were analyzed using high-resolution full-scan (SIM) acquisition over an m/z range of 70-1,030 at a resolution of 140,000, with an AGC target of 3×10^6^ and a maximum injection time of 500ms. Pooled quality control samples were additionally analyzed using data-dependent MS/MS (dd-MS^2^) to facilitate metabolite identification. For dd-MS^2^ acquisitions, full MS scans were acquired over an m/z range of 70-1,030 at a resolution of 70,000, with an AGC target of 1×10^6^ and a maximum injection time of 200ms. MS/MS spectra were acquired at a resolution of 17,500 with an AGC target of 1×10^5^, a maximum injection time of 50ms, a loop count of 15, an isolation window of 1.2m/z, and stepped normalized collision energies of 20 and 50.

#### Metabolomic Data Processing and Statistical Analysis

Raw data were processed using Compound Discoverer version 3.4 (Thermo Fisher Scientific). Metabolites were annotated through accurate mass and MS/MS spectral matching against the ChemSpider and mzCloud databases. Chromatographic peak areas were used for relative quantification and statistical comparison between experimental groups. A series of analytical standards representing a broad range of metabolites were analyzed using the same LC-MS/MS method and data processing workflow. These standards can be utilized to provide additional confidence in metabolite annotations generated by Compound Discoverer by aligning their retention times with corresponding features detected in the experimental samples. Prior to statistical analysis, normalized abundance tables were first consolidated to remove redundant annotations, di-, tri-, and tetra-peptides and metabolites not present in mammalian systems. Compounds were first filtered by identified matches in the HMDB All metabolites (v5. 217719 cpds) column (part of the raw data processing workflow). Where a single compound was represented by multiple entries, the one with the highest mean abundance across samples and with the expected retention time was retained. One liver sample (LTreated2) was excluded before analysis because of abnormally low tissue weight. The remaining samples in each tissue were screened for outliers using three criteria, computed on log2-transformed metabolite intensities: 1. Within-group correlation: the mean Pearson correlation of each sample with the other samples in its group, computed leaving that sample out. A sample was flagged if this fell below the median minus three scaled median absolute deviations across all samples of that tissue. 2. Silhouette width: computed on per-metabolite z-scored data, using Euclidean distance with experimental groups as clusters. A sample was flagged if its silhouette width was below 0. 3. Robust Mahalanobis distance: calculated from a minimum covariance determinant (MCD) estimate on the first three principal components of the z-scored data. A sample was flagged if its robust distance exceeded 3.06, the square root of the 97.5th percentile of the chi-square distribution with 3 degrees of freedom. Samples flagged by all three criteria were excluded. One liver sample (LCre6) met the criterion; no heart samples did. The final analysis therefore included 18 heart samples and 16 liver samples (5 WT, 6 Unrecombined, 5 Recombined). Normalized metabolite abundances were log_2_-transformed, with non-positive values (below detection) set to missing. For each metabolite, differences among the three groups were assessed by one-way ANOVA, followed by Tukey’s honestly significant difference (HSD) post-hoc test for the three pairwise contrasts; group means and log_2_ fold changes were computed on the log_2_-transformed values. ANOVA and Tukey p-values were corrected for multiple testing across all metabolites using the BH FDR procedure, applied independently to the omnibus test and pairwise differences were considered significant at Tukey p_adj_<0.05. After analysis, significant metabolites were manually screened for any abnormal annotations (those that should not be present in mammalian systems) that bypassed initial filtering and were subsequently removed from the list. Analyses were performed in R 4.6.1 using base functions (aov, TukeyHSD, p.adjust, robustbase, cluster). Normalized abundances and analyzed data are available in **Suppl. File 3**.

### NBTx Staining

Frozen OCT blocks containing liver, intestine, gastrocnemius muscle, spleen, or heart tissue were sectioned onto slides using a Leica CM 1860 Cryostat in 10µm sections and stored at −80°C until they were ready to be stained. NBTx staining protocol was adapted from Simard et al.(*18*). Briefly, sections were circled using a PAP pen, fully covered with NBTx solution and incubated at 16°C for 5 minutes (heart), 12 minutes (intestine), 30 minutes (muscle), 1 hour (spleen) or 15 minutes (liver). Slides were then rinsed in PBS and dehydrated in 70% ethanol, 95% ethanol, 100% ethanol (2x) and xylene before being coverslipped with Cytoseal 60. Slides were imaged on an Echo Revolve microscope at 4x, 10x, and 20x magnifications. Images were analyzed and scored for their % NBTx positive area (or positive cells per 100 micron for liver) using FIJI ImageJ software.

### Bulk RNA-Sequencing

Heart and liver tissue from 16 month old, male mice (n=6/group) was dounced in Trizol/Chloroform and homogenized with zirconia beads for 20 minutes at 4°C. Samples were then centrifuged at 10,000g for 5 minutes and supernatant purified according to the RiboPure RNA Purification Kit (Invitrogen, cat#: AM1924). RNA was DNase treated and samples were sent to Novogene for library preparation using their standard pipeline and sequenced on an Illumina NovaSeq X Plus instrument. Trimmed paired-end reads were quantified with Kallisto quant against a custom Mus musculus transcriptome generated from Ensembl release 116 (GRCm39), restricted to GENCODE primary protein-coding and lncRNA transcripts. Transcript abundance estimates were summed by gene symbol and analyzed separately for each tissue in R using DESeq2 (version 1.40.2). Genes with zero counts in any sample were excluded. Unwanted variation was estimated using sva (version 3.48.0) and RUVSeq (version 1.34.0), using empirically identified invariant genes as controls. Differential expression among the three experimental groups was assessed using a likelihood-ratio test (LRT), and genes with a Benjamini–Hochberg-adjusted (p_adj_<0.05) were considered significant. Significant genes were clustered by expression pattern using DEGreport. Gene Ontology and Reactome pathway enrichment analyses were performed using clusterProfiler (version 4.8.2), with the genes tested by DESeq2 used as the background. Pairwise comparisons were also determined by DESeq2. Sequencing data are being deposited to SRA. Analyzed data are available in **Suppl. File 4**.

### Dorsal Fibroblast Isolation, Culture, and *in vitro* Experiments

WT, *Mfn1^loxp^*, *Polg*^D257A-flox^ and *Polg*^D257A-flox^; *Mfn1^loxp^* mice were euthanized at 2-months of age, and dorsal skin fibroblasts were isolated following the cited protocol(*34*). Fibroblasts were maintained in typical culture media (DMEM, 10% FBS, 1% P/S). When the cells initially filled up the entire culture plate, fibroblasts were split into three dishes (n=3/genotype). Since the *Polg*^D257A-flox^ and *Polg*^D257A-flox^; *Mfn1^loxp^* cells were to be used for mutation tracking, it is essential to note that although the three lines were derived from a single animal, these mice were unrecombined at their euthanasia; therefore, to ensure a diverse mutational spectrum and for the three lines to essentially become biological replicates in their mutational profiles, we passaged unrecombined cells for three weeks prior to *in vitro* recombination, so that each line would have ample time to create its own mutational profile.

### *In vitro* recombination of *Polg* and *Mfn1* alleles

Each cell line (n=3/genotype) was recombined *in vitro* by overnight incubation with Cre Recombinase Gesicles (Takara Bio, #631449) to recombine both the *Polg* and *Mfn1* alleles. After 48 hours, *Polg* recombination was verified as above (see **Cre Recombination Strategies** above) and *Mfn1* knockout was validated by Western blot (see **Western Blot** below) and by mitochondrial staining with MitoTracker Deep Red FM (Thermo Fisher) and subsequent imaging at 63x magnification on a Leica Stellaris confocal microscope. Refer to **DNA extraction & Duplex-Sequencing** above for those methods.

### Pearson Syndrome (PS) Cybrid Cell Lines, *MFN1* Knockout and Heteroplasmy Measurement

PS1 90% heteroplasmy cybrid 143B cells were obtained from the laboratory of Sonia Emperador at the University of Zaragoza(*35*) and were grown in standard cell culture media, supplemented with 50µg/mL uridine. DMEM without penicillin-streptomycin was used when performing transfections and transductions. Cells were subsequently transduced to stably express Cas9 endonuclease (ABM # K002, Origene #TR30037). Knockout of *MFN1*: Cas9-expressing PS1 90% cells were transfected with a plasmid containing two gRNAs for *MFN1* (gRNA1: GUUAUAUGGCCAAUCCCACU and gRNA2: GAUGAUCUGGUAGAAAUGCA) or a plasmid containing two scramble gRNAs and with Lipofectamine 3000 Transfection Reagent (Thermo Fisher). Plasmids were generated by VectorBuilder (VB240903). Cells underwent antibiotic selection. Subsequently, monoclonal populations of successfully transfected cells were generated through limiting dilution cloning, and knockout of *MFN1* was verified by live cell imaging of mitochondria for punctate appearance with Mitotracker Green FM (Thermo Fisher) on a Leica Stellaris confocal microscope at 63x magnification. Measurement of deletion burden: DNA was extracted as described above, and used to measure relative mitochondrial DNA heteroplasmy by qPCR (Bio-Rad CFX Connect Real-Time PCR System) using one primer pair that spans the common deletion to measure deletion presence (F: 5’-CCCCCATACTCCTTACACTATTCC-3’ and R: 5’-TGCGGTTTCGATGATGTGGT-3’) and one primer pair to measure total number of mtDNA molecules in the 12S rRNA (F: 5’-AGCGCAAGTACCCACGTAAA-3’, R: 5’-TGCTAAATCCACCTTCGACCC-3’) with SYBR Green iTaq Supermix (Bio-Rad cat#: 1725124).

### Western Blot (Tissues & Cell Lines)

Whole cell lysates were generated by homogenizing cells or tissue in radioimmunoprecipitation buffer (RIPA). Total protein was determined by BCA assay (Thermo Fisher). 20µg of protein were boiled at 75°C under denatured conditions and resolved on 4–20% gradient gels. Proteins were electroblotted using a Criterion blotter (Bio-Rad Laboratories) and transferred onto 0.45μm polyvinyl difluoride membranes. Membranes were stained using Revert 700 fluorescent protein stain and imaged before blocking with LI-COR Intercept blocking buffer (LI-COR Biosciences), followed by primary antibody incubation overnight at 4°C (anti-MFN1, abcam ab221661). Membranes incubated with IRDye 800CW and/or 700CW secondary antibodies and visualized by Odyssey (LI-COR Biosciences). Western blot data were quantified with ImageJ and normalized by total protein per lane.

### Immunofluorescence Staining

Frozen OCT blocks containing heart or liver tissue were sectioned onto slides using a Leica CM 1860 Cryostat in 10µm sections and stored at −80°C until they were ready to be stained. Slides were air-dried for 10 minutes. For cleaved caspase-3 (Cell Signaling #9661) and TNF (Cell Signaling #11948), slides were fixed and permeabilized with 100% acetone for 10 minutes at −20°C. For Ki67 (Abcam #ab15580), slides were fixed with 4% PFA for 10 minutes at room temperature, washed 3x with PBS, and permeabilized with 0.02% Digitonin for 10 min at room temperature. All slides were then washed 3x with PBS. Sections were blocked using blocking buffer (0.1% Tween-20, 2.5% BSA, 5% Normal Goat Serum in PBS) for 1 hour at room temperature. Sections were then incubated with primary antibody at 1:150 dilution in a humidified chamber overnight at 4°C. Slides were washed in PBS 3×5min, and then sections were incubated with secondary antibody Goat anti-Rabbit Alexa Fluor 488 (Invitrogen, #A-11034) at 1:800 dilution in the dark for 1 hour at room temperature. Slides were washed in PBS 3×5min, mounted using Vectashield Antifade Mounting Medium with DAPI (Vector Labs, #H-1200-10), and coverslipped. Slides were imaged on the Leica Stellaris Confocal microscope at 63x magnification. All images were analyzed using ImageJ. All images were quantified by positive cell count over total number of nuclei.

### Statistics

Group differences means (± SEM) were analyzed by one-way ANOVA with Tukey’s test, Kruskal-Wallis with Dunn’s test for nonparametric distributions, or Welch’s ANOVA when variances were unequal across groups. For comparisons between two groups, Welch’s t-test was performed. Significance was defined as p<0.05. Echocardiography, NBTx quantification, and immunofluorescence quantification were performed in a blinded manner and imaging analysis utilized threshold-based quantification in FIJI with identical settings applied across all groups. No animals or data were excluded except as specified in the relevant Methods sections. Plots were made and analyses were performed using GraphPad Prism version 11 (GraphPad Software, San Diego, CA).

---

**Supplemental Figure S1.**
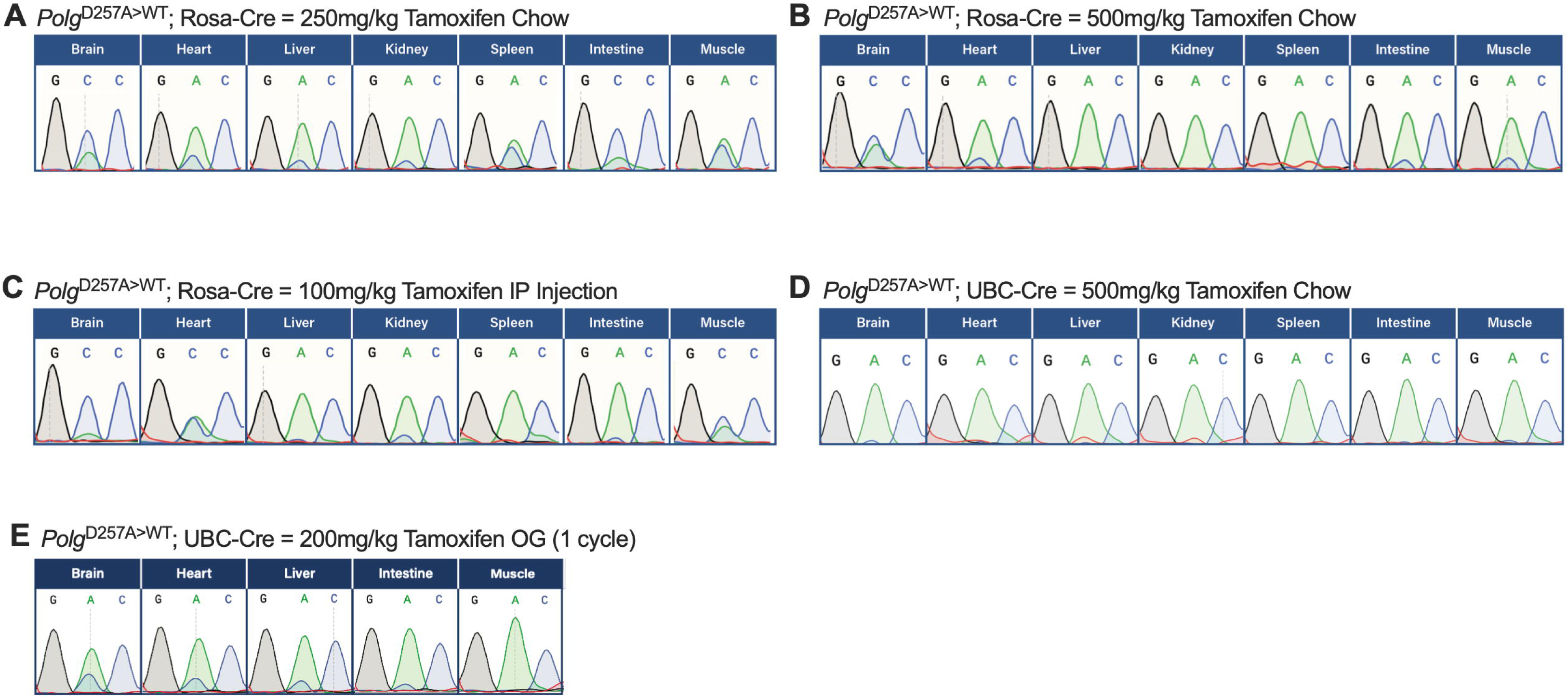
Sanger sequencing traces of cDNA to assess recombination efficiency of various tested strategies. (A-C) *Polg*^D257A-flox^ mice were initially crossed to ROSA26-CreER^T2^ mice and administered 250mg/kg tamoxifen chow *ad libitum* for 4 weeks **(A)**, 500mg/kg tamoxifen chow *ad libitum* for 4 weeks **(B)**, or 100mg/kg tamoxifen intraperitoneal injections once daily for 5 days **(C)**. Recombination efficiency was not sufficient to conduct the experiments proposed, as even low levels of ongoing mutagenesis would confound the study. **(D,E)** *Polg*^D257A-flox^ mice were then crossed to UBC-CreER^T2^ mice and administered 500mg/kg tamoxifen chow *ad libitum* for 4 weeks **(D)** or 200mg/kg tamoxifen by oral gavage once daily for 5 days **(E)**.

**Supplemental Figure S2.**
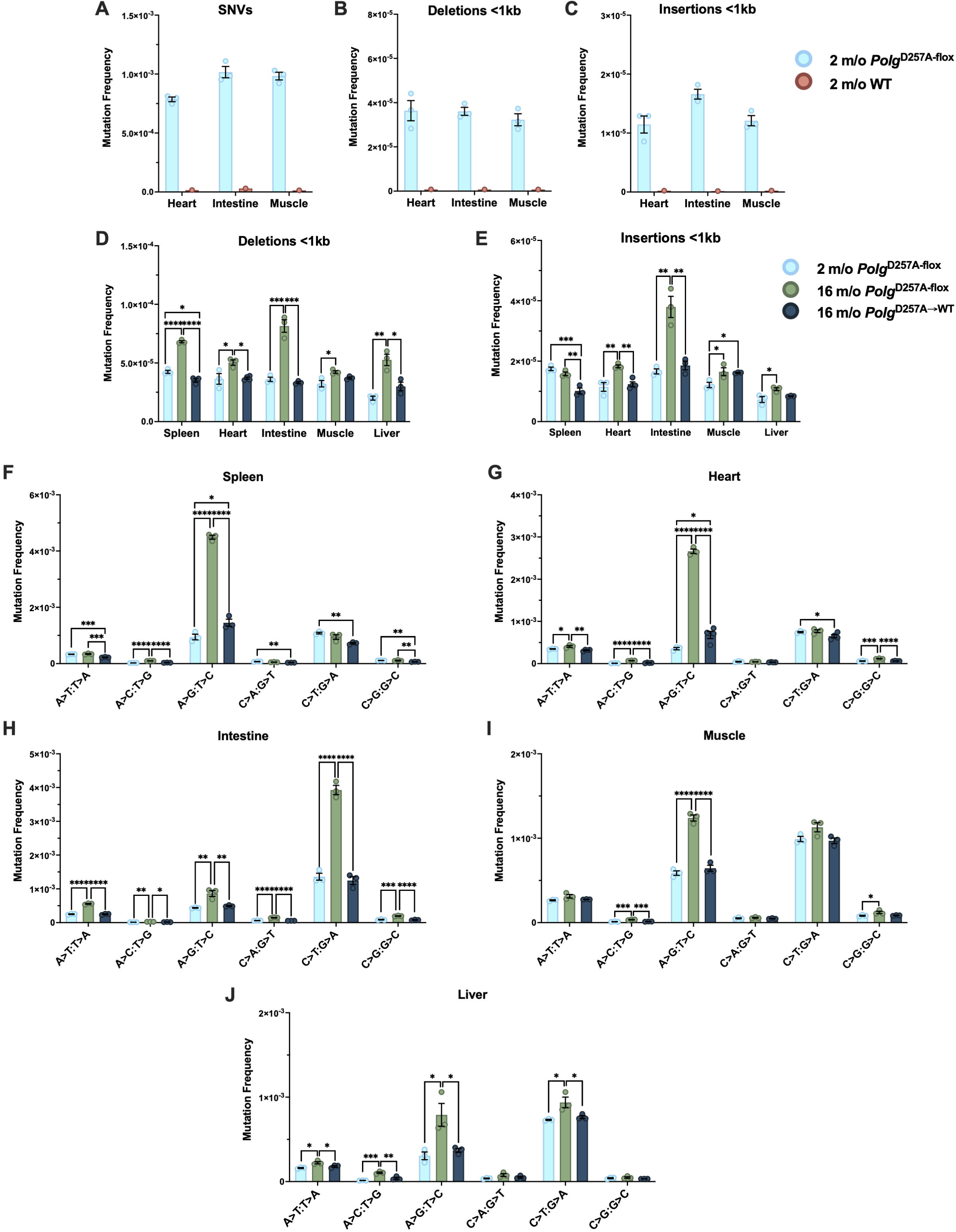
Mutational profiles of 2-month-old animals and mutation spectra of 16-month- old animals across tissues. (A-C) Mutation frequency determined by duplex sequencing of SNVs **(A)**, deletions **(B)**, and insertions **(C)** in heart, intestine, and muscle tissue from 2-month-old *Polg*^D257A-flox^ (n=3) and one WT mouse (n=1). **(D,E)** Deletion **(D)** and insertion **(E)** mutation frequency determined by duplex sequencing in spleen, heart, intestine, and muscle in aged experimental animals compared to 2-months. **(F-J)** Mutation spectra across spleen **(F)**, heart **(G)**, intestine **(H)**, muscle **(I)**, and liver **(J)**, in 2- and 16-month-old animals (n=3-4/group). Group differences means (± SEM) were analyzed by one-way ANOVA with Tukey’s test. *p < 0.0332, **p<0.0021, ***p<0.0002,****p<0.0001.

**Supplemental Figure S3.**
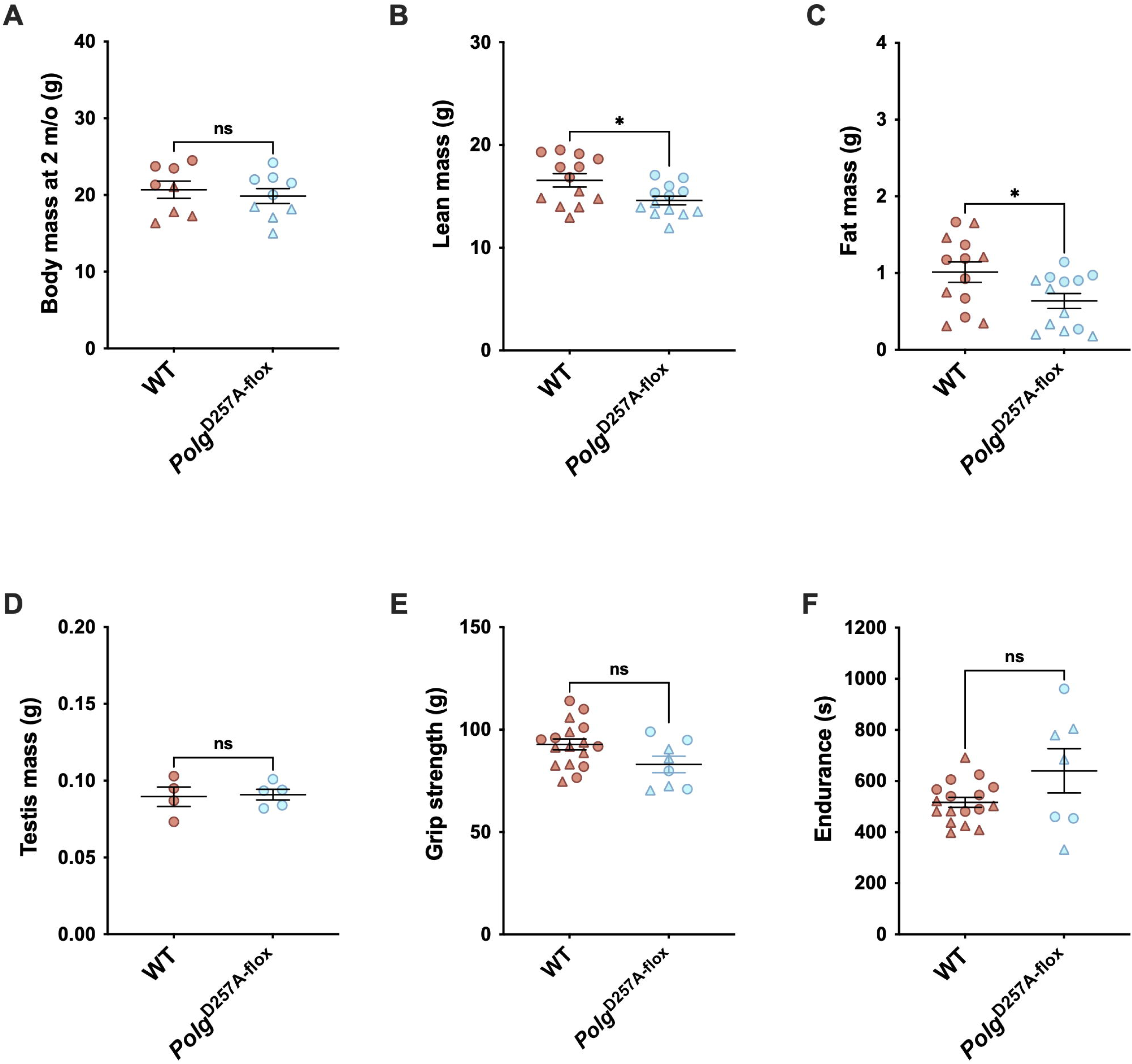
2-month-old physiology in WT and *Polg*^D257A-flox^ mice. **(A)** Body weights at 2-months of age (n=4-5/group/sex). **(B,C)** Body composition at 2-months of age, with lean mass in **(B)** and fat mass in **(C;** n=6-7/group/sex). **(D)** Testes are the same mass at 2-months in both WT and *Polg*^D257A-flox^ animals (n=4-5/group). **(E,F)** Grip strength **(E)** and endurance **(F)** are the same in WT and *Polg*^D257A-flox^ mice at 2-months-old (male n=4-9/group/sex). Group differences means (± SEM) were analyzed by Welch’s t-test. *p < 0.0332, **p<0.0021, ***p<0.0002,****p<0.0001. Males are denoted by: , while females: .

**Supplemental Figure S4.**
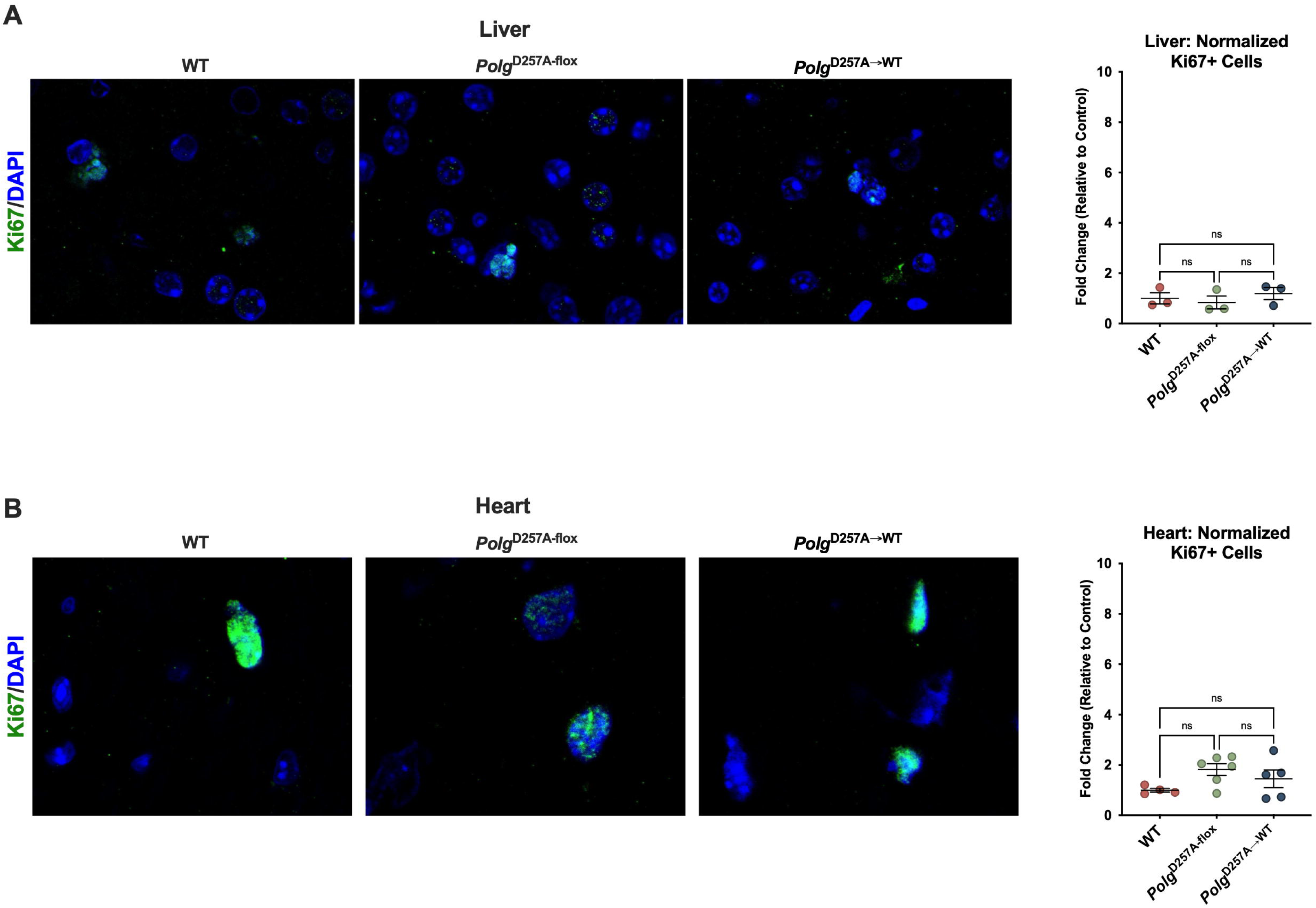
Ki67 immunofluorescence staining in 16-month old males in liver and heart tissue. (A,B) Liver staining **(A)** and quantification **(B)** represented by fold change relative to control (n=3/group). **(C,D)** Heart staining **(C)** and quantification **(D)** represented by fold change relative to control (n=4-6/group). Both tissues show no change across experimental groups in Ki67 positive cells. Images taken at 63x magnification. Group differences means (± SEM) were analyzed by one-way ANOVA with Tukey’s test. *p < 0.0332, **p<0.0021, ***p<0.0002,****p<0.0001.

**Supplemental Figure S5.**
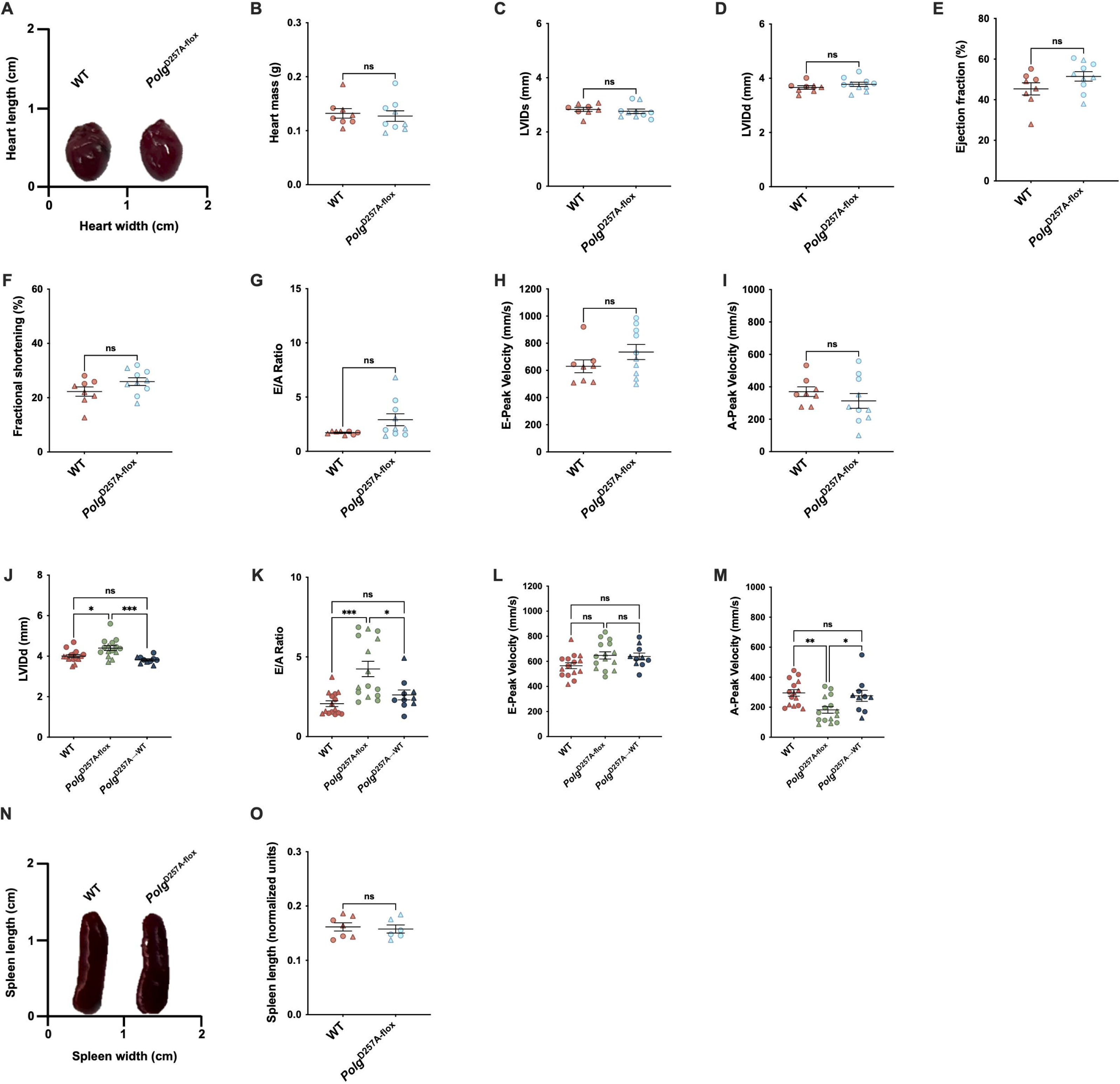
Heart and spleen phenotypes at 2-months of age, and additional 16-month-old cardiac phenotypes. (A-I) Hearts of 2-month-old WT and *Polg*^D257A-flox^ mice are the same in size **(A)**, mass **(B;** n=4-5/group/sex), and function, as determined by LVIDs **(C)**, LVIDd **(D)**, ejection fraction **(E)**, fractional shortening **(F)**, E/A Ratio **(G)**, E-peak velocity **(H)**, and A-peak velocity **(I;** male n=3-5/group; female n=5/group). **(J-M)** Diastolic function is impaired in 16-month-old *Polg*^D257A-flox^ mice, but not in WT or *Polg*^D257A→WT^ mice as demonstrated by the left ventricle internal diameter in diastole (LVIDd) **(J)**, E/A ratio **(K)**, E-peak velocity **(L)**, A-peak velocity **(M)**, highlighting specifically atrial filling dysfunction (male n=6-9/group; female n=4-7/group). **(N,O)** Spleens of WT and *Polg*^D257A-flox^ mice are also the same in overall size **(N)** and length **(O)** at 2-months of age (n=3-4/group/sex). Group differences means (± SEM) were analyzed by Welch’s t-test. *p < 0.0332, **p<0.0021, ***p<0.0002,****p<0.0001. Males are denoted by: , while females: .

**Supplemental Figure S6.**
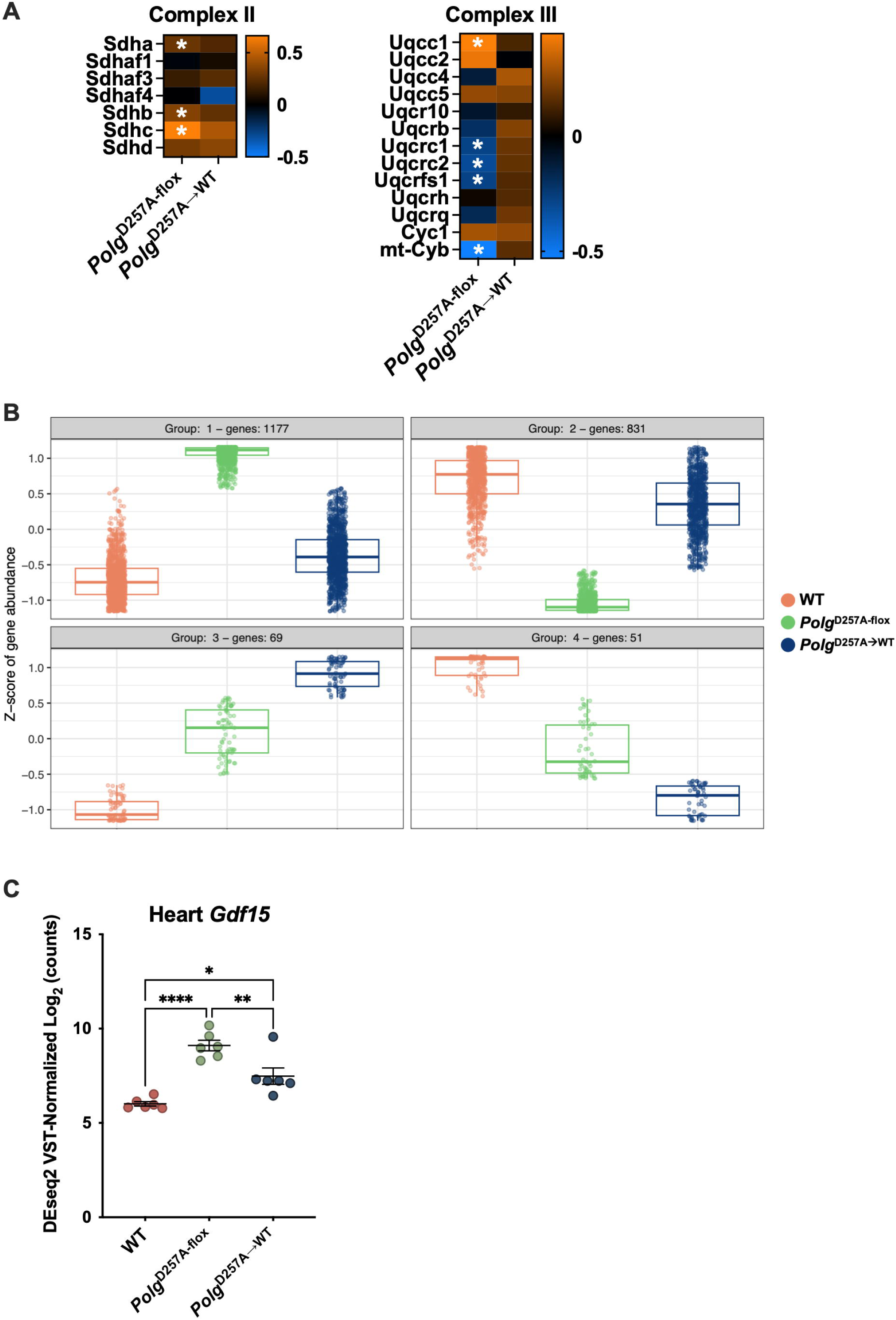
Multi-omic characterization of heart tissue across genotypes. **(A)** Heatmaps of differential protein expression in *Polg*^D257A-flox^ and *Polg*^D257A→WT^ hearts compared to WT of nuclear- and mitochondrial-encoded ETC Complexes II and III. Asterisks denote significantly altered proteins versus WT. **(B)** k-means clustering of differentially expressed genes (RNA-seq) in heart across the three genotypes. Each panel is one gene cluster, with the number of genes per cluster indicated above; the y-axis shows the z-scored gene abundance across samples, and boxplots show the median and interquartile range with individual genes as points. **(C)** VST-normalized log_2_ counts of *Gdf15* transcripts in the hearts of WT, *Polg*^D257A-flox^ and *Polg*^D257A→WT^ mice. Group differences means (± SEM) were analyzed by one-way ANOVA with Tukey’s test. *p < 0.0332, **p<0.0021, ****p<0.0001. Transcriptomics: n=6/genotype, proteomics: n=6/genotype, significance by BH p_adj_<0.05.

**Supplemental Figure S7.**
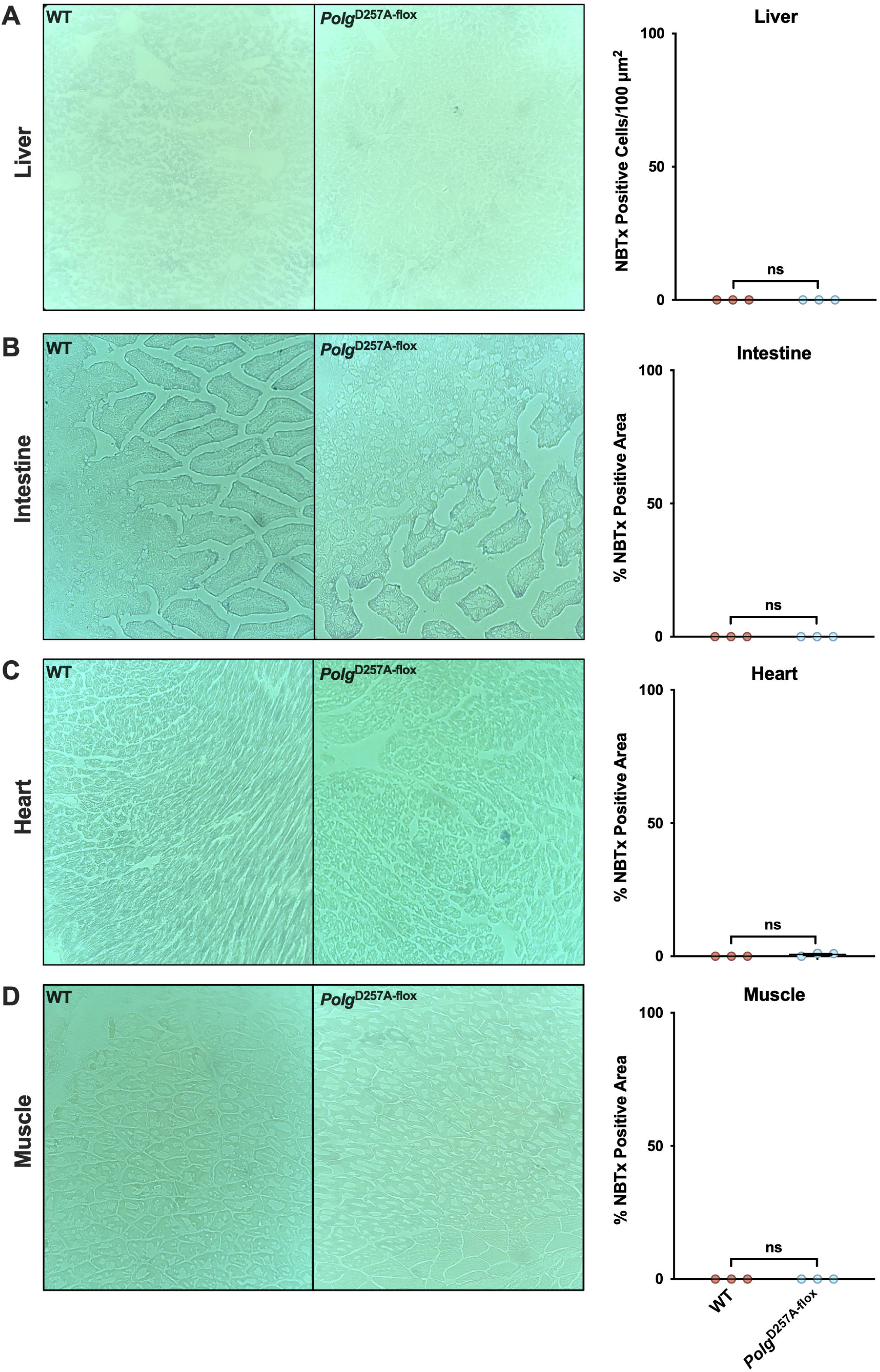
NBTx staining for clonal expansion in 2-month old animals. (A-D) NBTx staining and quantification of liver **(A)**, intestine **(B)**, heart **(C)**, and gastrocnemius muscle **(D)**. There are no clonally expanded mutations in any of the tissues at 2-months of age in WT and *Polg*^D257A-flox^ male mice (n=3/group). Images at 10x magnification.

**Supplemental Figure S8.**
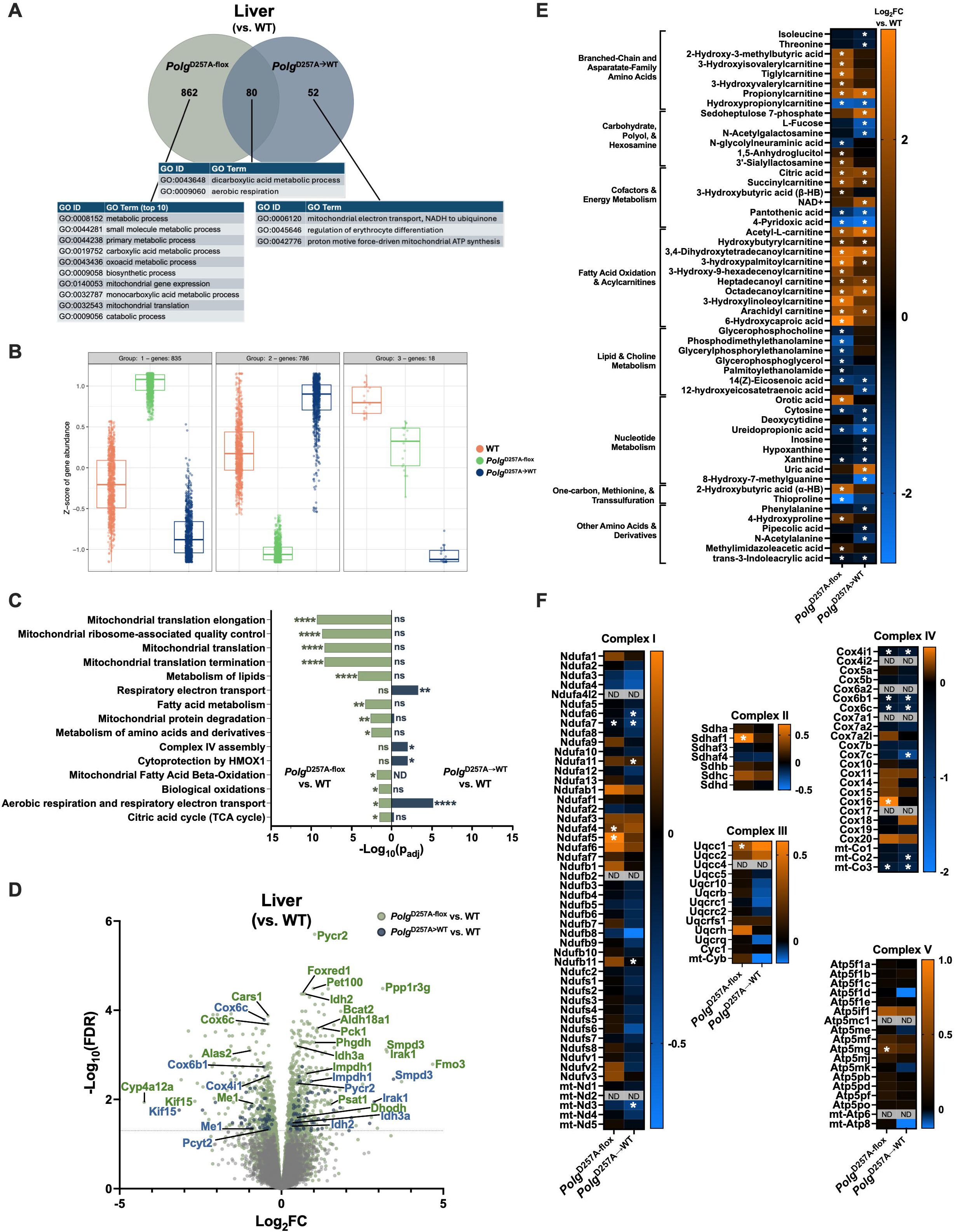
Early-life mutations leave a persistent molecular scar in liver tissue. **(A)** Unique and overlapping differentially expressed proteins between *Polg*^D257A-flox^ or *Polg*^D257A→WT^ and WT in liver. Top 10, or all, Gene Ontology (GO; biological process) terms are displayed in the table for significantly altered proteins. GO enrichment analysis was performed using the GO Knowledgebase, and pathways with padj<0.05 were considered significant. **(B)** k-means clustering of differentially expressed genes (RNA-seq) in liver across the three genotypes. Each panel is one gene cluster, with the number of genes per cluster indicated above; the y-axis shows the z-scored gene abundance across samples, and boxplots show the median and interquartile range with individual genes as points. **(C)** ORA pathways significantly enriched in *Polg*^D257A-flox^ and *Polg*^D257A→WT^ livers compared to WT, plotted by significance (−log_10_p_adj_). **(D)** Volcano plot of differentially expressed proteins for liver between *Polg*^D257A-flox^ and WT (green) or *Polg*^D257A→WT^ and WT (blue). Select proteins are labeled. **(E)** Heatmap of differential metabolite abundance between *Polg*^D257A-flox^ or *Polg*^D257A→WT^ and WT. **(F)** Heatmap of differential protein expression in *Polg*^D257A-flox^ and *Polg*^D257A→WT^ livers compared to WT of nuclear- and mitochondrial-encoded ETC Complexes. Asterisks denote significantly altered proteins vs. WT. ND = not detected. Transcriptomics: n=6/genotype, proteomics: n=6/genotype, significance by BH p_adj_<0.05; metabolomics n=5-6/genotype, significance by one-way ANOVA with Tukey’s post-hoc test p_adj_<0.05.

**Supplemental Figure S9.**
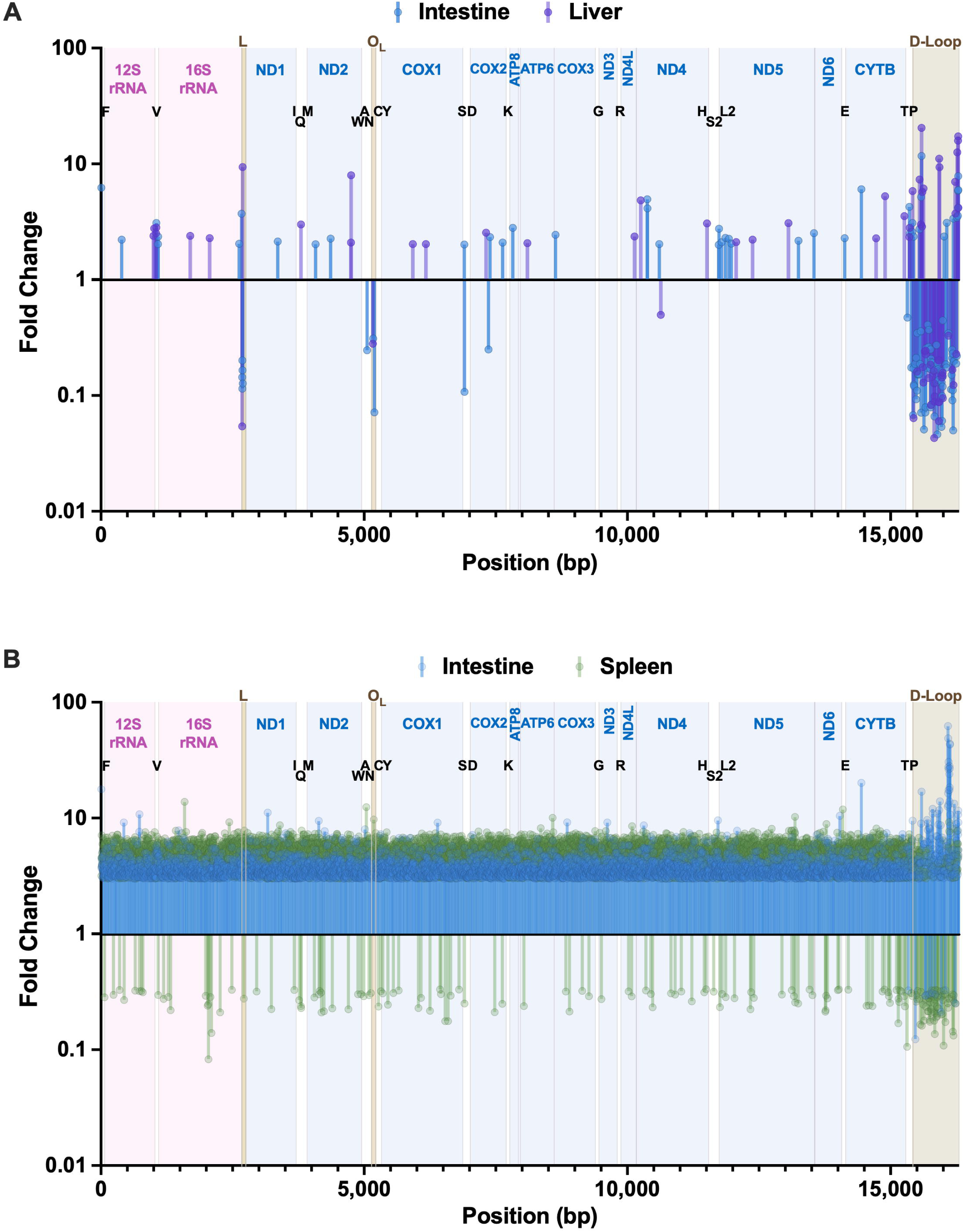
**Significantly altered mutations with age in *Polg*^D257A^**^→**WT**^ **and *Polg*^D257A-flox^ mice along the genome. (A)** Significant absolute >2-fold changes in mutation frequency over time in liver (purple) and intestine (blue) of *Polg*^D257A→WT^ mice. Mutations in regulatory regions are depleted, while mutations in protein coding genes are broadly either not significantly changed or increase in frequency over time, along the genome. **(B)** Significant absolute >3-fold changes in mutation frequency over time in spleen (green) and intestine (blue). Mutations overwhelmingly increase over time in these tissues in *Polg*^D257A-flox^ mice all along the genome. Significance by beta binomial testing, BH q<0.05; n=3/group.

**Supplemental Figure S10.**
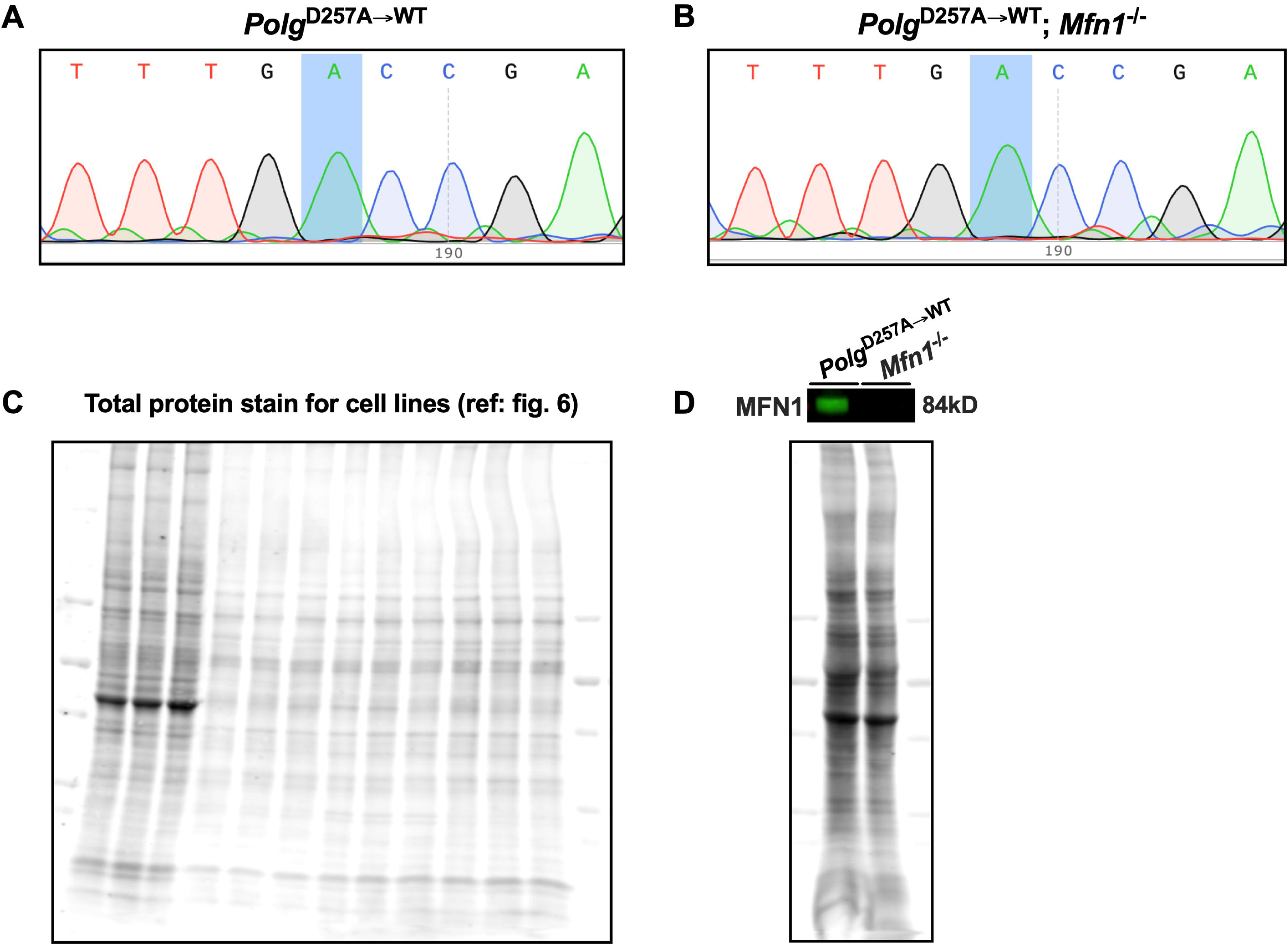
Supporting data for validation of *Polg* and *Mfn1* allelic recombination. (A,B) Sanger sequencing traces of *Polg* locus in *Polg*^D257A→WT^ (**A**) and *Polg*^D257A→WT^; *Mfn1*^−/-^ (**B;** mutant GCC *Polg* WT *Polg* GAC) cell lines after Cre Gesicle-mediated *in vitro* recombination. Both cell lines express WT *Polg* after recombination. **(C)** Total protein stain for western blot normalization of *Mfn1* protein expression from **Fig. 6** in cell lines (left, n=3/genotype). **(D)** Western blot of *Polg*^D257A→WT^ and *Polg*^D257A→WT^; *Mfn1*^−/-^ intestines, demonstrating knockout of MFN1 in this tissue (n=1/genotype shown).

**Supplemental Figure S11.**
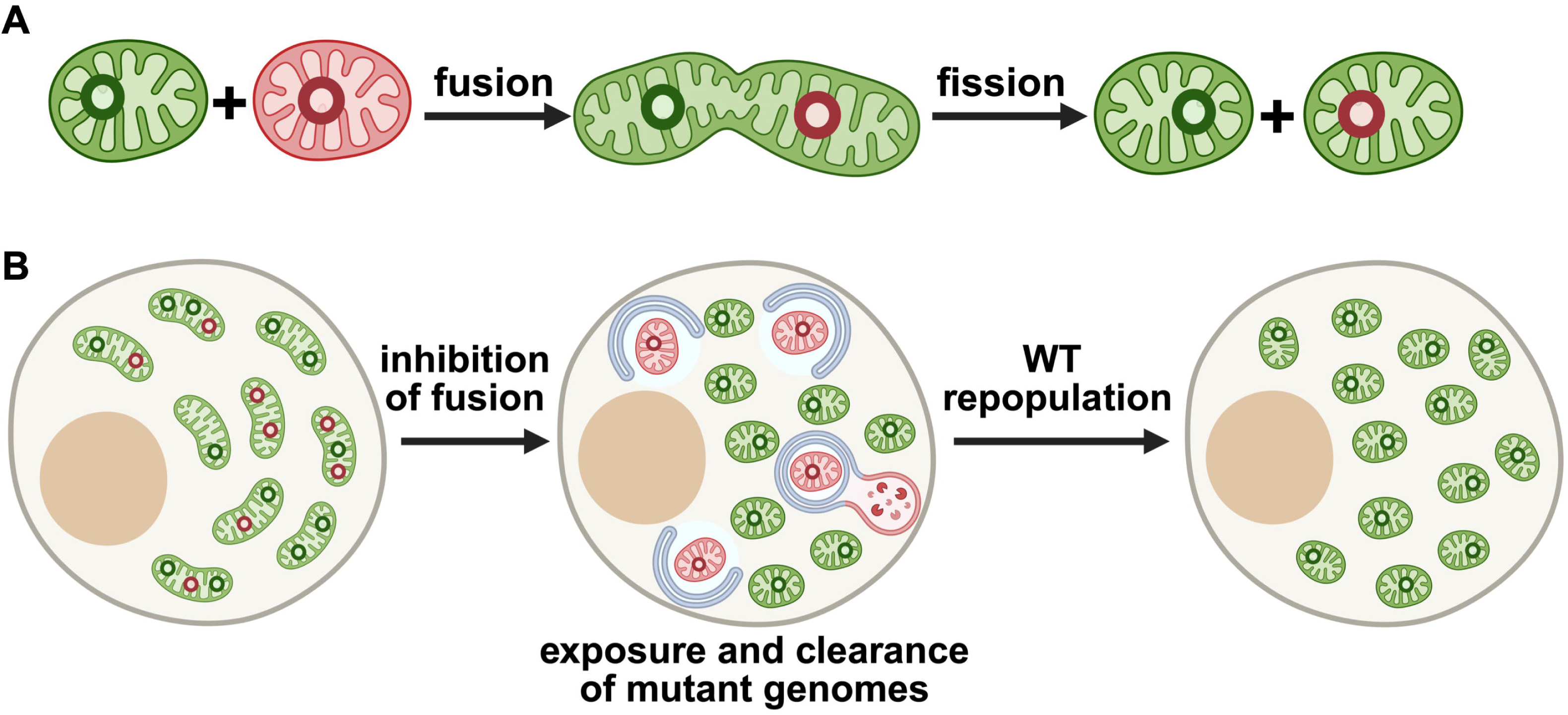
Schematic of model for the selective clearance of mutant genomes upon fusion inhibition. **(A)** Mitochondrial fusion allows dysfunctional mitochondria (red mitochondrion) carrying mutant genomes (red genome) to receive WT proteins, RNA, and mtDNA (green genome) from functional mitochondria (green mitochondrion) through complementation, rescuing their dysfunction. Subsequent fission yields functional mitochondria that nonetheless carry mutant genomes (green mitochondrion, red genome) **(B)** Although complementation through fusion allows cells to tolerate large mtDNA mutation burdens, it also masks the presence of mutant genomes inside of seemingly healthy organelles (left). Inhibition of fusion forces mitochondria to rely on their own genomes to sustain energy production, which results in their removal by mitophagy when they can no longer do so (center). WT genomes then repopulate the mitochondrial pool (right). Created with BioRender.

## REFERENCES

1. D. C. Wallace, Mitochondrial DNA mutations in disease and aging. Environ Mol Mutagen 51, 440–450.

2. K. Ishikawa, K. Takenaga, M. Akimoto, N. Koshikawa, A. Yamaguchi, H. Imanishi, K. Nakada, Y. Honma, J. Hayashi, ROS-generating mitochondrial DNA mutations can regulate tumor cell metastasis. Science 320, 661–664 (2008).

3. A. A. Elorza, J. P. Soffia, mtDNA Heteroplasmy at the Core of Aging-Associated Heart Failure. An Integrative View of OXPHOS and Mitochondrial Life Cycle in Cardiac Mitochondrial Physiology. Front Cell Dev Biol 9, 625020 (2021).

4. T. Nomiyama, Y. Tanaka, N. Hattori, K. Nishimaki, K. Nagasaka, R. Kawamori, S. Ohta, Accumulation of somatic mutation in mitochondrial DNA extracted from peripheral blood cells in diabetic patients. Diabetologia 45, 1577–1583 (2002).

5. S. W. Ballinger, J. M. Shoffner, E. V. Hedaya, I. Trounce, M. A. Polak, D. A. Koontz, D. C. Wallace, Maternally transmitted diabetes and deafness associated with a 10.4 kb mitochondrial DNA deletion. Nature genetics 1, 11–15 (1992).

6. Y. Kraytsberg, E. Kudryavtseva, A. C. McKee, C. Geula, N. W. Kowall, K. Khrapko, Mitochondrial DNA deletions are abundant and cause functional impairment in aged human substantia nigra neurons. Nature genetics 38, 518–520 (2006).

7. A. W. Linnane, S. Marzuki, T. Ozawa, M. Tanaka, Mitochondrial DNA mutations as an important contributor to ageing and degenerative diseases. Lancet 1, 642–645 (1989).

8. C. Lawless, L. Greaves, A. K. Reeve, D. M. Turnbull, A. E. Vincent, The rise and rise of mitochondrial DNA mutations. Open Biol 10, 200061 (2020).

9. L. C. Greaves, M. Nooteboom, J. L. Elson, H. A. Tuppen, G. A. Taylor, D. M. Commane, R. P. Arasaradnam, K. Khrapko, R. W. Taylor, T. B. Kirkwood, J. C. Mathers, D. M. Turnbull, Clonal expansion of early to mid-life mitochondrial DNA point mutations drives mitochondrial dysfunction during human ageing. PLoS genetics 10, e1004620 (2014).

10. A. Bolden, G. P. Noy, A. Weissbach, DNA polymerase of mitochondria is a gamma-polymerase. The Journal of biological chemistry 252, 3351–3356 (1977).

11. G. C. Kujoth, A. Hiona, T. D. Pugh, S. Someya, K. Panzer, S. E. Wohlgemuth, T. Hofer, A. Y. Seo, R. Sullivan, W. A. Jobling, J. D. Morrow, H. Van Remmen, J. M. Sedivy, T. Yamasoba, M. Tanokura, R. Weindruch, C. Leeuwenburgh, T. A. Prolla, Mitochondrial DNA mutations, oxidative stress, and apoptosis in mammalian aging. Science 309, 481–484 (2005).

12. A. Trifunovic, A. Wredenberg, M. Falkenberg, J. N. Spelbrink, A. T. Rovio, C. E. Bruder, Y. M. Bohlooly, S. Gidlof, A. Oldfors, R. Wibom, J. Tornell, H. T. Jacobs, N. G. Larsson, Premature ageing in mice expressing defective mitochondrial DNA polymerase. Nature 429, 417–423 (2004).

13. S. R. Kennedy, M. W. Schmitt, E. J. Fox, B. F. Kohrn, J. J. Salk, E. H. Ahn, M. J. Prindle, K. J. Kuong, J. C. Shen, R. A. Risques, L. A. Loeb, Detecting ultralow-frequency mutations by Duplex Sequencing. Nature protocols 9, 2586–2606 (2014).

14. M. Vermulst, J. Wanagat, G. C. Kujoth, J. H. Bielas, P. S. Rabinovitch, T. A. Prolla, L. A. Loeb, DNA deletions and clonal mutations drive premature aging in mitochondrial mutator mice. Nature genetics 40, 392–394 (2008).

15. M. Vermulst, J. H. Bielas, G. C. Kujoth, W. C. Ladiges, P. S. Rabinovitch, T. A. Prolla, L. A. Loeb, Mitochondrial point mutations do not limit the natural lifespan of mice. Nature genetics 39, 540–543 (2007).

16. C. Trumpff, Q. Huang, J. Michelson, C. C. Liu, D. Shire, C. G. Habeck, Y. Stern, M. Picard, Blood mitochondrial health markers cf-mtDNA and GDF15 in human aging. bioRxiv, (2025).

17. M. Yang, K. H. Vousden, Serine and one-carbon metabolism in cancer. Nat Rev Cancer 16, 650–662 (2016).

18. M. L. Simard, A. Mourier, L. C. Greaves, R. W. Taylor, J. B. Stewart, A novel histochemistry assay to assess and quantify focal cytochrome c oxidase deficiency. J Pathol 245, 311–323 (2018).

19. W. McLaren, L. Gil, S. E. Hunt, H. S. Riat, G. R. Ritchie, A. Thormann, P. Flicek, F. Cunningham, The Ensembl Variant Effect Predictor. Genome Biol 17, 122 (2016).

20. M. Falkenberg, N. G. Larsson, C. M. Gustafsson, Replication and Transcription of Human Mitochondrial DNA. Annual review of biochemistry 93, 47–77 (2024).

21. E. Yakubovskaya, E. Mejia, J. Byrnes, E. Hambardjieva, M. Garcia-Diaz, Helix unwinding and base flipping enable human MTERF1 to terminate mitochondrial transcription. Cell 141, 982–993 (2010).

22. Y. Goto, I. Nonaka, S. Horai, A mutation in the tRNA(Leu)(UUR) gene associated with the MELAS subgroup of mitochondrial encephalomyopathies. Nature 348, 651–653 (1990).

23. T. Lieber, S. P. Jeedigunta, J. M. Palozzi, R. Lehmann, T. R. Hurd, Mitochondrial fragmentation drives selective removal of deleterious mtDNA in the germline. Nature 570, 380–384 (2019).

24. N. P. Kandul, T. Zhang, B. A. Hay, M. Guo, Selective removal of deletion-bearing mitochondrial DNA in heteroplasmic Drosophila. Nat Commun 7, 13100 (2016).

25. S. B. Hannah Tobias-Wallingford, Lauren Gaspar, Brian Gallagher, Carole Bassa, Emmanuel Sotirakis, Vincent Dubus, Christopher Hemme, Lars Olson, Giuseppe Coppotelli, Jaime M. Ross, A novel inducible mtDNA mutator mouse model to study mitochondrial dysfunction with temporal and spatial control. BioRxiv, (2025).

26. S. B. Ong, A. R. Hall, D. J. Hausenloy, Mitochondrial dynamics in cardiovascular health and disease. Antioxid Redox Signal 19, 400–414 (2013).

27. D. F. Dai, L. F. Santana, M. Vermulst, D. M. Tomazela, M. J. Emond, M. J. MacCoss, K. Gollahon, G. M. Martin, L. A. Loeb, W. C. Ladiges, P. S. Rabinovitch, Overexpression of catalase targeted to mitochondria attenuates murine cardiac aging. Circulation 119, 2789–2797 (2009).

28. G. C. Kujoth, P. C. Bradshaw, S. Haroon, T. A. Prolla, The role of mitochondrial DNA mutations in mammalian aging. PLoS genetics 3, e24 (2007).

29. A. Ventura, D. G. Kirsch, M. E. McLaughlin, D. A. Tuveson, J. Grimm, L. Lintault, J. Newman, E. E. Reczek, R. Weissleder, T. Jacks, Restoration of p53 function leads to tumour regression in vivo. Nature 445, 661–665 (2007).

30. Y. Ruzankina, C. Pinzon-Guzman, A. Asare, T. Ong, L. Pontano, G. Cotsarelis, V. P. Zediak, M. Velez, A. Bhandoola, E. J. Brown, Deletion of the developmentally essential gene ATR in adult mice leads to age-related phenotypes and stem cell loss. Cell Stem Cell 1, 113–126 (2007).

31. H. Chen, J. M. McCaffery, D. C. Chan, Mitochondrial fusion protects against neurodegeneration in the cerebellum. Cell 130, 548–562 (2007).

32. J. G. Hoekstra, M. J. Hipp, T. J. Montine, S. R. Kennedy, Mitochondrial DNA mutations increase in early stage Alzheimer disease and are inconsistent with oxidative damage. Ann Neurol 80, 301–306 (2016).

33. M. Sanchez-Contreras, M. T. Sweetwyne, B. F. Kohrn, K. A. Tsantilas, M. J. Hipp, E. K. Schmidt, J. Fredrickson, J. A. Whitson, M. D. Campbell, P. S. Rabinovitch, D. J. Marcinek, S. R. Kennedy, A replication-linked mutational gradient drives somatic mutation accumulation and influences germline polymorphisms and genome composition in mitochondrial DNA. Nucleic Acids Res 49, 11103–11118 (2021).

34. A. Seluanov, A. Vaidya, V. Gorbunova, Establishing primary adult fibroblast cultures from rodents. Journal of visualized experiments : JoVE, (2010).

35. C. Hernandez-Ainsa, E. Lopez-Gallardo, M. C. Garcia-Jimenez, F. J. Climent-Alcala, C. Rodriguez-Vigil, M. Garcia Fernandez de Villalta, R. Artuch, J. Montoya, E. Ruiz-Pesini, S. Emperador, Development and characterization of cell models harbouring mtDNA deletions for in vitro study of Pearson syndrome. Dis Model Mech 15, (2022).

